# Discovery of cell active macrocyclic peptides with on-target inhibition of KRAS signaling

**DOI:** 10.1101/2021.09.11.459913

**Authors:** Shuhui Lim, Nicolas Boyer, Nicole Boo, Chunhui Huang, Gireedhar Venkatachalam, Yu-Chi Angela Juang, Michael Garrigou, Kristal Kaan, Ruchia Duggal, Khong Ming Peh, Ahmad Sadruddin, Pooja Gopal, Tsz Ying Yuen, Simon Ng, Srinivasaraghavan Kannan, Christopher J. Brown, Chandra Verma, Peter Orth, Andrea Peier, Lan Ge, Xiang Yu, Bhavana Bhatt, Feifei Chen, Erjia Wang, Nianyu Jason Li, Raymond J. Gonzales, Alexander Stoeck, Brian Henry, Tomi K. Sawyer, David Lane, Charles W. Johannes, Kaustav Biswas, Anthony W. Partridge

**Author notes:** These authors contributed equally to this work (Shuhui Lim, Nicolas Boyer).

## Abstract

RAS is the most commonly mutated oncogene in human cancers and RAS-driven tumors are amongst the most difficult to treat. Historically, discovery of therapeutics targeting this protein has proven challenging due to a lack of deep hydrophobic pockets to which a small molecule might bind. The single such pocket available is normally occupied by GDP or GTP which have millimolar cellular concentrations and picomolar affinities for KRAS and hence is challenging to target. The recent FDA approval of sotorasib, a covalent modifier of the KRAS^G12C^ mutant protein, has clinically validated KRAS as an oncology target. However, traditional challenges remain for targeting the more common KRAS mutations such as G12D and G12V. As an alternative approach, we investigated the superior binding capacity of macrocyclic peptides to identify KRAS inhibitory molecules. We focused on the recently reported disulfide-cyclized arginine-rich peptide **KRpep-2d**, discovered through phage display and previously independently confirmed by us as a *bona fide* KRAS binder. To mitigate intracellular disulfide reduction and loss of binding, we identified an alternate cyclization motif by inverting the configuration of Cys5 and linking it to Cys15 through a thioacetal bridge. The corresponding peptide bound KRAS through cis isomerization of the peptide bond between D-Cys5 and Pro6 as observed through x-ray crystallography. Prototypic molecules displayed measurable cellular inhibition of RAS signaling without membrane lysis and counter-screen off-target activity. An analogue containing azido-lysine confirmed the cell penetrant nature of this molecule using our recently reported NanoClick assay. To improve cellular activity, we sought to improve proteolytic stability. Structure-activity relationship studies identified key scaffold residues critical for binding and revealed that replacement of N- and C-terminal arginine residues with D-arginines and introduction of an α-methyl moiety at Ser10 resulted in a molecule with improved proteolytic stability as indicated by its persistence in whole cell homogenate. The resulting peptide **MP-3995** had improved and sustained cell-based efficacy. On-target activity was validated through confirmation of target engagement, lack of signaling in irrelevant pathways, and use of non-binding control peptides. Mechanism of action studies suggested that peptide binding to both GDP- and GTP-states of KRAS may contribute to cellular activity. Although validated as *bona fide* and on-target inhibitors of cell based KRAS signaling, this series is unlikely to advance to the clinic in its current form due to its arginine-dependent cell entry mechanism. Indeed, we showed a strong correlation between net positive charge and histamine release in an *ex vivo* assay making this series challenging to study *in vivo*. Nonetheless, these molecules provide valuable templates for further medicinal chemistry efforts aimed at leveraging this unique inhibitory binding site on KRAS.

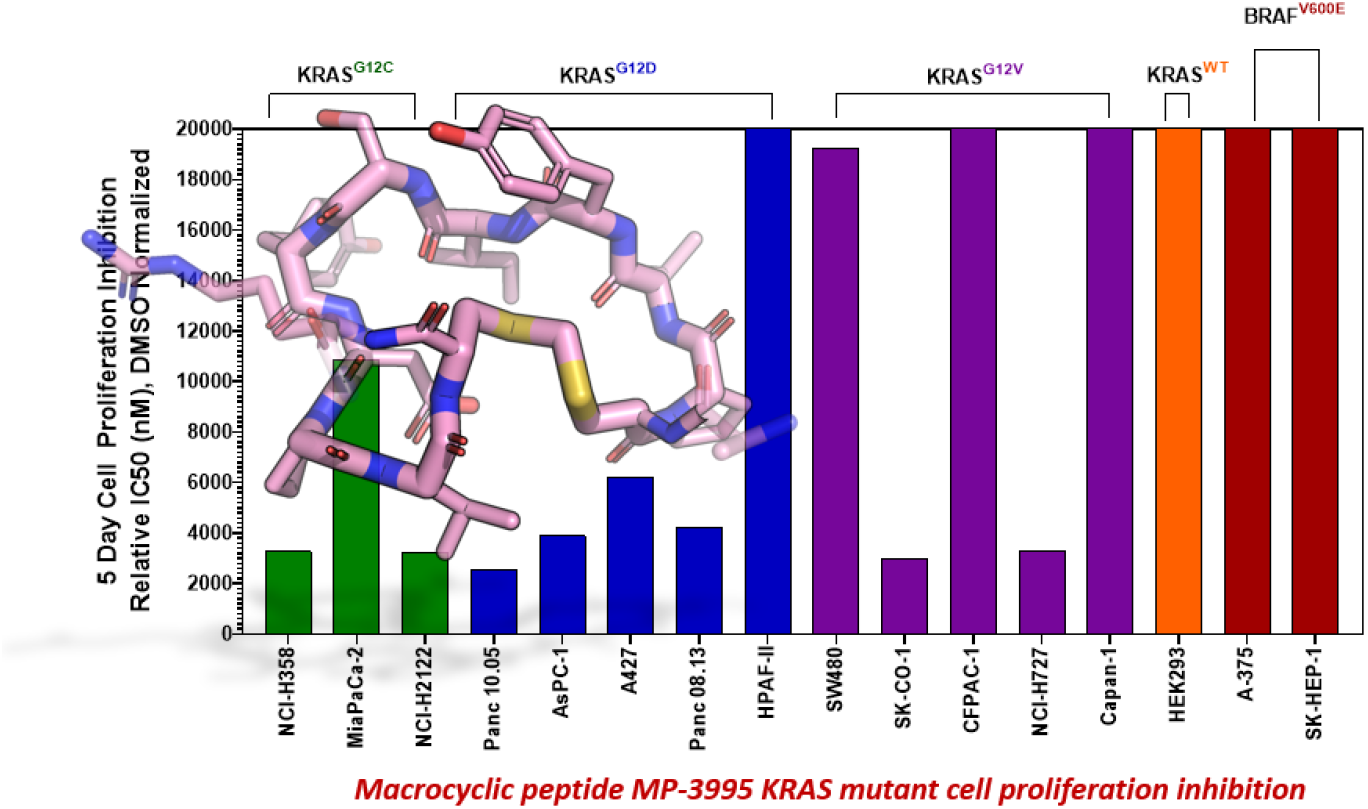

## Introduction

The RAS GTPase serves as a molecular switch to activate signaling cascades related to cell survival and proliferation, most notably, the MAPK and AKT pathways. Human malignancies gain growth advantages by mutating RAS at positions G12, G13, or Q61 to bias the protein to the signaling-active GTP-loaded state [1]. Indeed, RAS is the most mutated oncogene across human cancers. Amongst the different isoforms, KRAS is the most frequently mutated and is especially prevalent in pancreatic, lung, and colorectal cancers [1]. Although mutant KRAS was discovered to be a common driver of human cancers in the early 1980s, there were, until very recently, no approved therapeutics against this target. Fortunately, recent advances have reinvigorated the field. Small molecule covalent inhibitors of KRAS^G12C^ have shown efficacy in animal models and in the clinic [2, 3]. One of these, sotorasib (AMG 510, LUMAKRAS−), was recently approved for treatment in patients with KRAS^G12C^ driven non-small cell lung cancers that are either metastatic or locally advanced [4]. Despite these remarkable advances, significant challenges remain for targeting tumors driven by KRAS with non-G12C mutations. Specifically, the current clinical molecules rely on a covalent modifier strategy that has strict specificity for the C12 residue, which is present in humans at a prevalence of only 3.4% in colorectal cancers and in 7.4% of non–small-cell lung cancer (NSCLC) cases [5]. Furthermore, as clinical studies progress with G12C inhibitors, resistance mechanisms to these new drugs are being mapped, including identification of additional changes to the KRAS protein itself [6]. For the remaining more prevalent KRAS mutations (e.g. G12D, G12V), significant challenges remain. Specifically, campaigns to identify small molecule binders to KRAS have largely failed due to a paucity of surface pockets suitable for small molecule docking. One available druggable pocket appears to be the nucleotide binding site, but it is normally occupied by GDP or GTP, which have millimolar cellular concentrations and picomolar affinities for KRAS, hence posing a major challenge to the development of competitive inhibitors. These challenges have prompted investigators to consider alternative approaches. Amongst these, peptide-based modulators hold promise due to their propensity to bind to diverse protein epitopes and modulate their activity. Advances in display-based technologies [7, 8] have led to successful screens on multiple protein targets resulting in novel peptide hits, increasing the promise of this emerging modality as a complementary way to modulate protein function.

Several research groups have indeed reported high affinity *bona fide* peptide binders to KRAS that represent valuable starting points for drug discovery [9-12]. In particular, Sakamoto *et al*. used phage display to identify a disulfide cyclized peptide with the sequence Ac-Arg1-Arg2-Arg3-Arg4-Cys5-Pro6-Leu7-Tyr8-Ile9-Ser10-Tyr11-Asp12-Pro13-Val14-Cys15-Arg16-Arg17-Arg18-Arg19-NH2 [10]. This molecule, termed **KRpep-2d**, bound KRAS^G12D^ with nanomolar affinity in both the GDP and GTP analog loaded states. Alanine-scanning mutagenesis identified Leu7, Ile9, and Asp12 as critical binding residues [13]. X-ray crystallography revealed the binding site to be near Switch II, allosterically blocking the interaction of KRAS with the guanine nucleotide exchange factor, SOS1 [14]. Subsequently we [15] and others [12] verified this peptide as having high-affinity and stoichiometric binding to KRAS. Although **KRpep-2d** represents a promising and novel KRAS binder, we concluded that structural modifications were required to render it cell-active. In particular, the disulfide crosslink is not expected to remain intact within the reducing environment of the cytosol. Furthermore, additional medicinal chemistry optimization could address potential peptide stability and permeability deficiencies.

To achieve cell permeability, cationic and hydrophobic residues are often incorporated into peptides targeting intracellular proteins. However, these design features can confound the interpretation of biochemical and functional assays [15]. Indeed, some recently reported putative peptidic KRAS inhibitors incorporating these elements were, in fact, false positives [15]. Thus, we took extra caution in our studies with the **KRpep-2d** scaffold since it contains several hydrophobic residues and a total of eight extracyclic arginines. In particular, we applied rigorous controls in the experiments described herein to ensure that the macrocyclic peptides we designed were not only *bona fide* KRAS binders but also authentic, on-target cell active molecules. These peptides therefore represent a new approach towards blocking KRAS-driven signaling beyond the G12C mutation, an area of high unmet need.

Using **KRpep-2d** as the starting point, we sought to improve binding affinity, increase proteolytic stability, and impart membrane permeability to advance a molecule capable of blocking cellular KRAS signaling. After exploring different strategies, we successfully replaced the disulfide bond with a thioacetal crosslink involving a D-Cys residue at position 5. X-ray crystallography revealed that peptides using this macrocyclization motif bind to KRAS in a similar manner but with a *cis* peptide bond between D-Cys and Pro6. Replacing the N- and C-terminal arginine residues with their D-amino acid counterparts produced a peptide with weak cellular activity. Introduction of an α-methyl group at Ser10 resulted in a molecule (**MP-3995**) with prolonged proteolytic stability and cellular blockade of pERK activity in KRAS^G12D^ (AsPC-1) cells but was inactive against a KRAS^WT^ cancer line (A375) harboring BRAF^V600E^, a MAPK pathway activating mutation that is downstream of KRAS. On-target cellular activity was further verified using non-binding peptide controls, counter-screens, and target engagement assays. In a panel of cancer cell lines, **MP-3995** inhibited proliferation in KRAS dependent lines but not in KRAS independent lines. Despite these favorable attributes, we identified the arginine-rich nature of this series to be a barrier for further development as strong histamine release was observed in an *ex vivo* assay. Initial attempts to reduce the flanking Arg residues resulted in a loss of membrane permeability and cellular activity. Nevertheless, the peptides described in this report represent a valuable scaffold for the genesis of novel validated inhibitors of KRAS signaling in cancer patients unserved by current successes with covalent G12C inhibition.

## Results

### Replacement of the KRpep-2d disulfide bridge with a D-Cys5–Cys15 thioacetal linkage results in a redox-stable, high affinity peptide

Sakamoto *et al*. [10] used phage display to identify the first example of a macrocyclic peptide with *bona fide* high-affinity binding to KRAS (**KRpep-2d**). The specific binding of this peptide to KRAS^G12D^ was previously independently validated by us using a suite of biophysical approaches including isothermal titration calorimetry (ITC), surface plasmon resonance (SPR), thermal shift assay (TSA), and hydrogen-deuterium exchange mass spectrometry (HDX-MS) [15]. Although this molecule provides an excellent starting point for medicinal chemistry efforts, it contains a disulfide linkage between Cys5 and Cys15, which might not remain intact in the reducing cytosolic environment and thus lose binding. Indeed, although SPR analysis confirmed high affinity binding (1.7 nM) for KRAS^G12D^ with the disulfide intact, binding was completely lost in the presence of DTT, a result presumably due to disulfide reduction (Fig 1A). Furthermore, a linear derivative of **KRpep-2d** where the two cysteines were replaced by L-serine, a cyclization incompetent isostere, exhibited no binding (data not shown). For **KRpep-2d**, redox sensitivity likely also contributed to the lack of cell-based pERK inhibition in our hands, as measured by Western blot (Fig S1). Accordingly, we sought to identify a peptide that maintained KRAS binding affinity but with a redox insensitive cross-link. Using a template with an *N*-terminal 6-azido-L-lysine residue, to enable cellular permeability measurements with the NanoClick assay [16], we explored a series of disulfide bridge replacements (Table 1). To assess the peptide binding affinities in high-throughput format, we used a TR-FRET assay employing GDP-loaded KRAS^G12D^ and a FAM-labeled **KRpep-2d** family member tracer peptide (see Supporting Information). This assay reports relative binding values whose EC_50_’s are right-shifted compared to the corresponding binding constants (K_D_’s) due to the high concentration of tracer peptide used. When the disulfide was replaced with a lactam moiety linking L-Asp at position 5 and L-Dap ((*S*)-2,3-diaminopropionic acid) at position 15, peptide **1**, no binding was detected. Intriguingly, the corresponding D-Asp5/L-Dap15 analog (**2**) exhibited modest biochemical activity (EC_50_ = 19331 nM) suggesting a potential benefit from incorporating a D-amino acid at position 5. Next, we explored thioacetal cross-links to connect the cysteine thiols. The L-Cys5/L-Cys15 thioacetal peptide (**3**) lost activity when compared to the parent disulfide (TR-FRET EC_50_ = 13740 nM), similar to previous findings [13]. Encouragingly, the D-Cys5/L-Cys15 thioacetal (**MP-6483**) had much improved biochemical potency (172 nM), a 11-fold improvement over **KRpep-2d** (Table 1). However, adding another atom to the linker in the D-Cys5/L-homoCys15 analog (**4**) led to a marked reduction in TR-FRET EC_50_ (4719 nM). Importantly, the most active peptide from this set of analogs, **MP-6483**, maintained its binding capacity in both oxidizing and reducing environments (Table 1, Fig S2), as assessed by TR-FRET. In agreement with the SPR results, **KRpep-2d** lost the capacity to bind to GDP-loaded KRAS^G12D^ in the presence of 1mM DTT (Table 1, Fig S2). **MP-6483** also showed a weak capacity to block pERK and pAKT signaling downstream of KRAS in the cellular context, (Figs 1B, C) with an absence of membrane disruption, as assessed by a LDH release assay (Fig 1D). However, **MP-6483**’s potency appeared to weaken significantly over time in AsPC-1 cells (a KRAS^G12D^ pancreatic cancer cell line), especially when assessed by the alphaLISA assay (Fig 1C), suggesting potential proteolytic stability issues.

**Figure 1:**
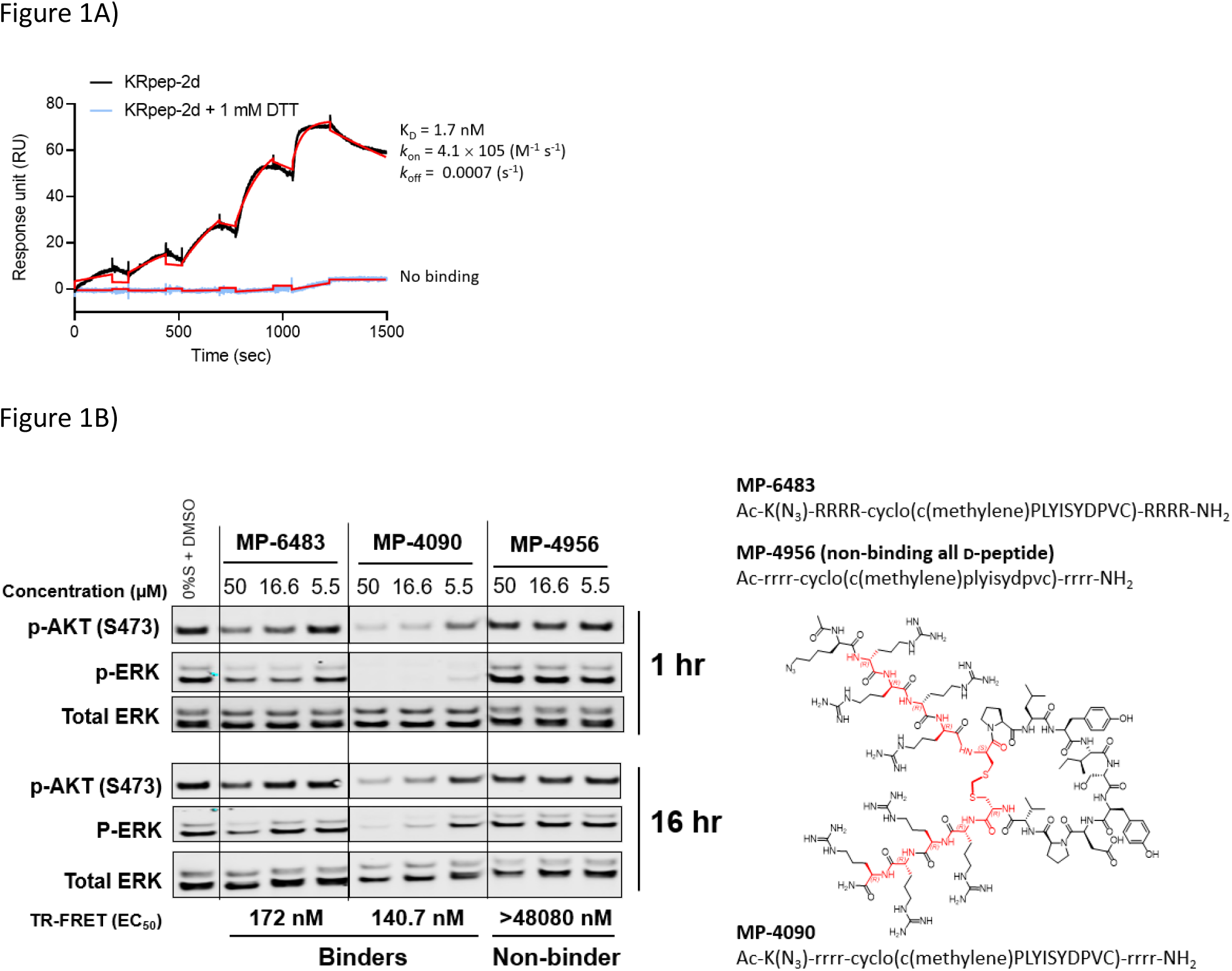
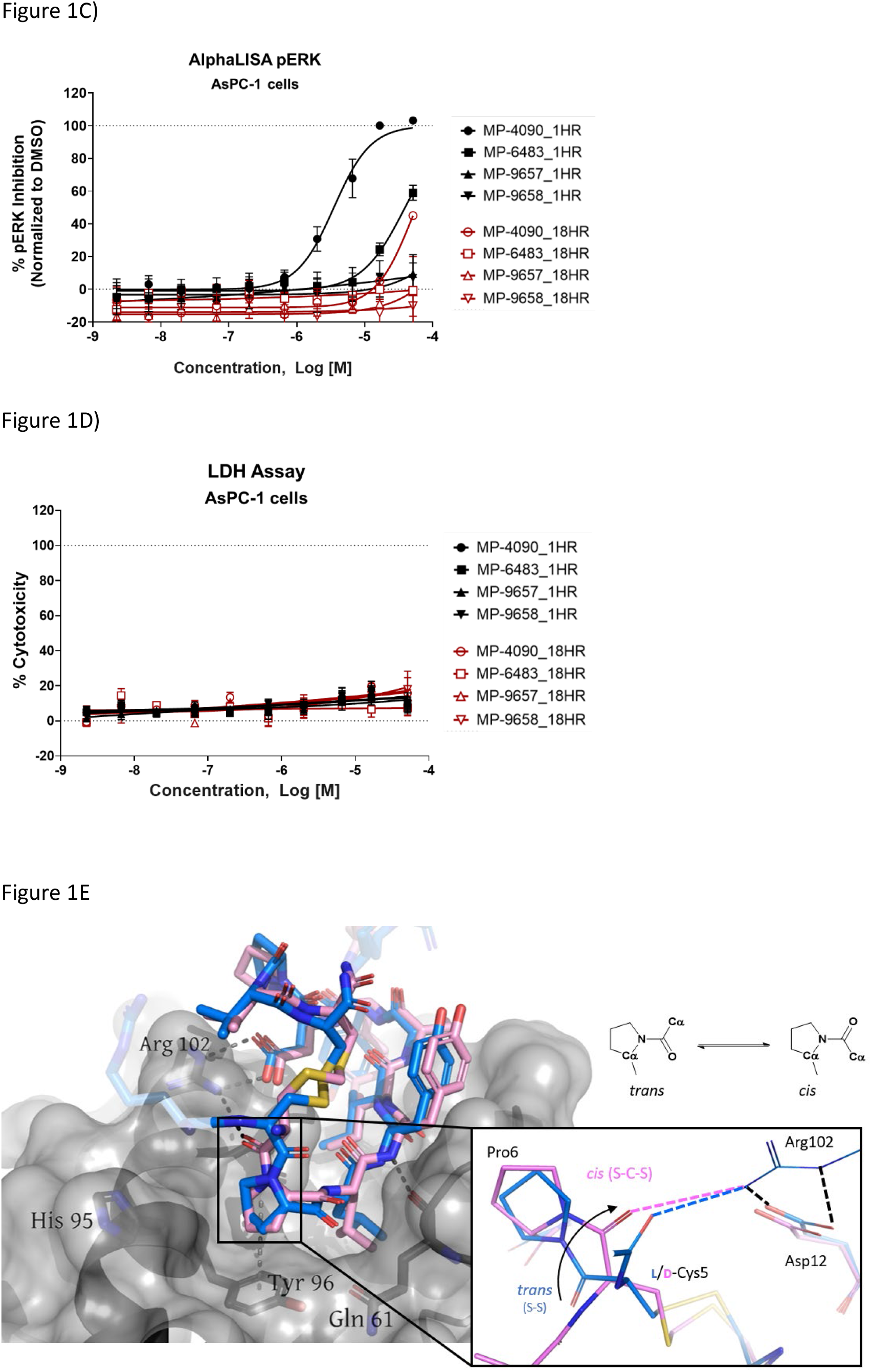
Replacement of the **KRpep-2d**’s disulfide bridge with a D-Cys5–CH_2_–L-Cys15 thioacetal linkage results in a redox-stable, high affinity peptide. A) SPR analysis shows that oxidized (cyclic) but not reduced (linear) **KRpep-2d** binds with high affinity to GDP-loaded KRAS^G12D^. B) Redox-stable peptides **MP-6483** and **MP-4090** display cell-based inhibition of KRAS signaling (pERK and pAKT) in AsPC-1 cells, as assessed by Western blot at 1-hour and 16-hour time-points; non-binding control **MP-4956** showed no activity (Left). Sequences for the peptides are shown (Top Right), along with the chemical structure of **MP-4090** (bottom right), the portions highlighted in red correspond to the exocyclic D-Arg backbone atoms and those from the thioacetal linker. C) **MP-6483** and **MP-4090** but not the non-binder controls (**MP-9657** and **MP-9658**) display cell-based inhibition (pERK) in AsPC-1 cells, as assessed by Alpha SureFire Ultra Multiplex Phospho/Total ERK1/2 assay (Perkin Elmer) when treated for 1 hr (n=6, black symbols) and 18 hrs (n=2, red symbols). D) The same lysates in panel C were also assessed for membrane toxicity, as measured by the CytoTox-ONE™ homogenous membrane integrity assay (Promega). E) Superimposition of co-crystal structures involving i) **KRpep-2d** in complex with KRAS^G12D^ (GDP) (PDB ID 5XCO, blue) and ii) **MP-9903**, a peptide containing the D-Cys5–CH_2_–L-Cys15 thioacetal linkage, with KRAS^G12D^ (GMPPCP) (PDB ID 7ROV, pink), shows a highly similar KRAS conformation and that binding of **MP-9903** involves a *cis* peptide bond between D-Cys5 and Pro6. For clarity, terminal arginines are either transparent or hidden.

**Table 1.** Modification of the disulfide cyclization motif of **KRpep-2d**

| Compound | Linker | KRAS<sup>G12D</sup> GDP TR-FRET EC<sub>50</sub> (nM) |
| --- | --- | --- |
| **KRpep-2d** | L-Cys5–L-Cys15 | 1916 (>50000<sup>a</sup>) |
| **1** | L-Asp5–L-Dap15 (lactam) | >50000 |
| **2** | D-Asp5–L-Dap15 (lactam) | 19331 |
| **3** | L-Cys5–CH<sub>2</sub>–L-Cys15 (thioacetal) | 13740 |
| **MP-6483** | D-Cys5–CH<sub>2</sub>–L-Cys15 (thioacetal) | 172 (93<sup>a</sup>) |
| **4** | D-Cys5–CH<sub>2</sub>–L-homoCys15 (thioacetal) | 4719 |

<sup>a</sup>Reducing conditions (1 mM DTT), see Figure S2.

### X-ray crystallography reveals a *cis* peptide bond conformation at D-Cys5–Pro6

To investigate the preference for the D configuration at position 5, we solved the co-crystal structure of KRAS loaded with a non-hydrolysable GTP analog (GMPPCP) bound to **MP-9903**, a peptide containing **MP-6483**’s macrocyclic sequence but with a reduced number of arginines at the N- and C-termini (see Table 4).

**KRpep-2d**’s structure was reported earlier in complex with KRAS GDP (PDB-ID 5XCO). Both structures overlap with a rms deviation of 0.51 Å excluding the switch I region, which undergoes conformational changes due to different crystal contacts (Fig 1E). Despite GMPPCP loading of KRAS in the **MP-9903** co-crystal structure, the protein adopts the off-state (GDP-loaded) conformation and the peptide binding pockets can easily be superimposed. Interestingly, the **MP-9903** peptide bond connecting D-Cys5 and Pro6 adopts a *cis* conformation allowing the cyclic part of the peptide to occupy the same binding site. Due to the opposite chirality at residue 5, the N-terminus has a more solvent exposed trajectory. Consequently, the hydrogen bond between the backbone carbonyl oxygen (O) of Arg4 and side chain nitrogen of KRAS/Arg102Nη is disrupted and Arg102 hydrogen bonds to the backbone carbonyl oxygen (O) of D-Cys5 due to the amide bond flip. All the other interactions observed between KRAS and **KRpep-2d** were also observed in the crystal structure of the KRAS – **MP-9903** complex (except the interactions that involve the terminal arginine) (Fig 1E). These interactions were also preserved during molecular dynamics simulations (Fig S3). The extension of the linker between D/L-Cys5 and Cys15 by a methylene (thioacetal) group increases the Cα–Cα distance by 0.65Å and remains solvent exposed. Similarly, the C-terminus of **MP-9903** is solvent exposed and lacks any interactions with KRAS. It matches the **KRpep-2d** conformation remarkably well (Fig 1E).

### Ala-scanning mutagenesis identifies key binding residues

To systematically address the binding specificity of the new macrocycle, we made a library of singular alanine substitutions focusing on the macrocyclic amino acids of **MP-1687**, a tetra-arginine variant of **MP-6483**. The resulting nine alanine mutants were tested for binding in the TR-FRET assay (Table S1). Four alanine mutants (Pro6Ala, Leu7Ala, Ile9Ala, Asp12Ala) caused a significant impairment of binding. Apart from the effect at the Pro6 position, these observations agree with those reported for the disulfide-bridged parent **KRpep-2d**, thus confirming the critical binding roles of Leu7, Ile9, and Asp12. Indeed, these results can be rationalized as the side chains of Leu7 and Ile9 are buried in the hydrophobic pocket on the surface of KRAS (Fig S3) and the side chain of Asp12 is involved in a salt bridge with the side chain of Arg102 from KRAS. Previous results in the context of **KRpep-2d** showed that Pro6Ala had only a modest (10-fold) reduction in affinity [13]. On the other hand, in the context of the extended thioacetal cross-link and inverted stereocenter at position 5, replacement of the constrained proline with an acyclic amino acid (Ala or *N*-Me-Ala) or modification of the ring size (azetidine-2-carboxylic acid (Aze) or pipecolic acid (Pip)) dramatically affected the bound conformation, thus resulting in dramatic loss of affinity (data not shown). This observation is in accordance with the *trans–cis* proline isomerization observed in the co-crystal structure and further suggested that local stabilization of the *cis* amide bond conformation might improve affinity. The side chains of residues Ser10 and Tyr11 interact with the side chain and backbone of KRAS residue Asp69 (Fig S3), thus rationalizing the moderate loss in affinity when these residues were substituted with alanine. Binding affinities for three other Ala mutants (i.e. Tyr8Ala, Pro13Ala, Val14Ala) were perturbed to lesser extent as the side chains of these residues are not involved in any intra or inter KRAS – peptide interactions (Fig S3). An *in silico* alanine scan was carried out using the crystal structure with standard protocols [17], and the results were found to be in reasonable accordance with experimental data; the smallest destabilization of +0.5 kcal/mol was seen for Val14Ala, while for the destabilized mutants (Pro6Ala, Leu7Ala, Ile9Ala, Asp12Ala) the calculations yielded values from 4-9 kcal/mol (Table S2).

### Replacing the terminal Arg residues with D-Arg results in a membrane permeable peptide with improved intracellular stability and on-target cellular activity

After identifying a binding competent, redox-stable thioacetal linkage, we then sought to improve peptide bioactivity by identifying metabolic soft spots and designing analogs with improved stability. Metabolite identification (MetID) studies on **KRpep-2d** were performed after incubation in homogenized THP1 cells and analysis via high resolution mass spectrometry. Rapid metabolism was observed, and several metabolites were detected within 10 mins that were a result of sequential loss of arginines from one or both termini, as evidenced by changes in mass to charge ratios. Ring opening resulting from disulfide bridge cleavage was not detected, likely due to the non-reducing conditions in the homogenized cells. Accordingly, we decided to replace all eight L-arginine residues on **MP-6483** with their hyper-stable enantiomeric counterpart (D-Arg) (Fig 1B). The resulting peptide, **MP-4090**, showed permeability as measured by the NanoClick assay (4-hour EC_50_ = 251 nM, 18-hour EC_50_ = 35 nM, Table 3), a result mirrored in imaging studies using FAM-labeled counterparts (Fig S4). *Importantly*, ***MP-4090*** *also showed, for the first time, robust inhibition of pERK and pAKT in AsPC-1 cells* (Fig 1B, C).

We also designed a series of non-binding control peptides including an all-D version of **MP-6483** (**MP-4956**) and peptides with enantiomeric substitutions at critical binding residues, Ile9 to D-Ile (**MP-9657)** and Asp12 to D-Asp (**MP-9658**) (Table 2). None of these peptides showed any binding in the TR-FRET assay (EC_50_ > 48080 nM) nor a capacity to inhibit cellular function (Fig 1B, C), despite evidence of permeability in the NanoClick assay (Table 2). They also did not induce membrane damage as assessed by the LDH release assay (Fig 1D). Furthermore, **MP-4090** and its non-binding control (**MP-9658**) were inactive in A375 and SK-MEL-28 cells (Fig S5A), counter-screen cell lines harboring BRAF^V600E^, a mutant that activates the MAPK pathway downstream of KRAS. AZ628 (BRAF inhibitor, [18]) and U0126 (MEK inhibitor, [19]), small molecules that inhibit the pathways downstream of KRAS, inhibited MAPK signaling in these BRAF mutant lines in the expected manner (Fig S5A). Live-cell imaging using FAM-labelled versions of these macrocyclic peptides confirmed that permeability was comparable between AsPC-1 and A375, but poorer in SK-MEL-28 (Fig S5B). In addition, **MP-4090** had no effect on upstream EGF receptor activation (Fig S6A) and TNFα-stimulated NFκB signaling (a KRAS-independent pathway) (Fig S6B) in AsPC-1 cells. This series of control experiments suggested that the cellular KRAS-inhibitory activity measured for **MP-4090** via inhibition of ERK phosphorylation was indeed on-target.

**Table 2.** Key peptides and non-binder controls on X-rrrr-cyclo(c(methylene)PLYISYDPVC)-rrrr-NH_2_ scaffold (**MP-4090**)

| Compound | X | Alteration | KRAS<sup>G12D</sup><br>GDP TR-FRET EC<sub>50</sub><br>(nM) | AsPC-1<br>pERK EC<sub>50</sub><br>1h/18h<br>(μM) | AsPC-1<br>LDH EC<sub>50</sub><br>1h/18h<br>(μM) | A375 pERK<br>EC<sub>50</sub> 1h/18h<br>(μM) | Nanoclick<br>EC<sub>50</sub> 4h/18h<br>(nM) |
| --- | --- | --- | --- | --- | --- | --- | --- |
| **MP-4090** | Ac-<br>K(N<sub>3</sub>)- | none | 140.7 | 3.6/30.5 | >50/>50 | >50/>50 | 250.5/34.9 |
| **MP-3995** | Ac-<br>K(N<sub>3</sub>)- | Ser10 to<br>α-Me-Ser | 172.1 | 1.2/1.5 | >50/>50 | >50/>50 | 1777/32.1 |
| **MP-4956** | Ac- | All D-peptide | >48080 | >50/>50 | >50/>50 | Not tested<br>/>50 | Not tested |
| **MP-9657** | Ac-K(N<sub>3</sub>)- | Ile9 to D-Ile | >48080 | >50/>50 | >50/>50 | Not tested | 1756/36.2 |
| **MP-9658** | Ac-K(N<sub>3</sub>)- | Asp12 to D-Asp | >48080 | >50/>50 | >50/>50 | Not tested />50 | 196.1/31.4 |

### Stabilizing the Tyr8/Ile9 metabolic soft spot gives a peptide with sustained cellular activity

Having validated **MP-4090** as a cell permeable peptide with on-target pERK inhibition, we next sought to improve its observed cellular potency. Although the exocyclic arginines had been replaced by their D-enantiomers, it appeared that the peptide was still susceptible to proteolytic cleavage as observed by the loss of pERK activity at 18 hours vs 1 hour (Fig 1C). Indeed, MetID studies on the **KRpep-2d** scaffold without the arginines performed in homogenized THP1 cells had also revealed an additional proteolytic soft spot between Tyr8 and Ile9 of the macrocycle.

To stabilize the peptide against proteolysis, we synthesized analogs with Tyr8 and Ile9 modifications including α-methylation and backbone homologation, however all were inactive in the TR-FRET assay (data not shown). We then expanded the α-methyl scan to additional residues and tested the resulting peptides for binding and stability (Table 3). Due to challenges with detecting highly cationic 8-Arg peptides on MS instrumentation, we carried out this study on the related 4-Arg analog (**MP-1687**, Table 4). α-Methylation was tolerated at positions 10 (Ser, **8**) and 13 (Pro, **11**) without any change in binding affinity when compared to **MP-1687**, with a modest 4-fold loss observed at position 14 (Val, **12**). In contrast, potency loss was observed at all other positions, ranging from 30-fold at Tyr11 to 800-fold at positions 8 or 15. Position 5 and 9 were not investigated. Comparison of the HeLa cell homogenate half-lives of the α-methylated analogs to the parent **MP-1687** showed improvements ranging from >10-fold for position 8 to >4-fold for position 10 and >3-fold for position 13 modifications. Considering both the potency and stability data, the α-methyl-Ser10 mutation was selected for inclusion in future designs. Incorporating the α-methyl-Ser10 modification into our previous lead sequence (**MP-4090**) led to **MP-3995**, a peptide with sustained pERK inhibition with EC_50_ values of 1.2 and 1.5 µM at 1 and 18 hours, respectively (Table 2), eliminating the 8-fold cell potency loss seen with **MP-4090** at 18 h vs. 1 h. This peptide also did not induce membrane damage as assessed by the LDH release assay and did not inhibit pathway signaling in the RAS-independent A375 cell line with the BRAF^V600E^ mutation (Table 2).

**Table 3.** α-methyl amino acid scan biochemical assay and cell homogenate half-life data on peptide Ac-K(N_3_)-RR-cyclo(c(methylene)PLYISYDPVC)-RR-NH_2_ (**MP-1687**)

| Compound | Modification | KRAS<sup>G12D</sup> GDP TR-FRET EC<sub>50</sub> (nM) | HeLa t<sub>1/2</sub> (min) |
| --- | --- | --- | --- |
| **MP-1687** | None | 60 | 37 |
| **5** | α-Me-Pro6 | 10830 | 142 |
| **6** | α-Me-Leu7 | 11510 | 114 |
| **7** | α-Me-Phe8 | 46120 | >373 |
| **8** | α-Me-Ser10 | 62 | 154 |
| **9** | α-Me-Tyr11 | 1800 | 71 |
| **10** | α-Me-Asp12 | 3705 | 41 |
| **11** | α-Me-Pro13 | 67 | 134 |
| **12** | α-Me-Val14 | 222 | NA |
| **13** | α-Me-Cys15 | 48080 | 120 |

**Table 4.** Arginine-truncation and mast cell degranulation SAR based on **MP-6483**, Ac-K(N_3_)-RRRR-cyclo(c(methylene)PLYISYDPVC)-RRRR-NH_2_

| Compound | N-term Arg<br>(pos. 1-4) | C-term Arg<br>(Pos. 16-19) | KRAS<sup>G12D</sup><br>GDP TR-FRET EC<sub>50</sub><br>(nM) | AsPC-1 pERK<br>EC<sub>50</sub> 1h/18h<br>(μM) | # of<br>Arg | rMCD<br>histamine<br>release<br>threshold<br>(μM) | NanoClick<br>EC<sub>50</sub> 4h/18h<br>(nM) |
| --- | --- | --- | --- | --- | --- | --- | --- |
| **MP-4090** | r r r r | r r r r | 140.7 | 3.6/30.8 | 8 | $\geq 0.14$ | 251/35 |
| **MP-6483** | R R R R | R R R R | 172.4 | 15.9/>50 | 8 | $\geq 0.14$ | 2693/636 |
| **14** | – R R R | R R R – | 75.8 | >50/>50 | 6 | $\geq 0.14$ | 3314/>10000 |
| **MP-1687** | – – R R | R R – – | 59.4 | >50/>50 | 4 | $\geq 3.7$ | 4552/4711 |
| **MP-9903** | – – – R | R – – – | 138.6 | >50/>50 | 2 | >100 | 6939/6828 |
| **15** | – – – – | R – – – | 187.6 | >50/>50 | 1 | >100 | >10000/4462 |
| **16** | – – – – | – – – – | 409.5 | >50/>50 | 0 | >100 | >10000/5136 |

### MP-4090 and MP-3995 specifically engage with cellular KRAS to inhibit its interactions with RAF–RBD

Further evidence of the on-target nature of this peptide series was provided by the capacity of **MP-4090** and **MP-3995**, but not two non-binding controls (**MP-9657** and **MP-9658**), to specifically engage and thermally stabilize KRAS in an isothermal cellular thermal shift assay (CETSA) using intact AsPC-1 cells (Fig 2A). Target engagement was also supported by the capacity of **MP-4090** and **MP-3995** to inhibit the interaction of KRAS with an engineered binding reporter; a RAF RBD-CRD-eGFP fusion protein that was introduced into AsPC-1 cells using mRNA transfection. In the presence of DMSO or incubation with non-binder peptides **MP-9658** or **MP-9657**, there was a distinct accumulation of the eGFP fluorescence at the cell membrane, presumably due to its interaction with KRAS. However, when the experiment was repeated with the addition of **MP-4090** or **MP-3995**, this staining pattern was lost and the eGFP signal became diffuse (Fig 2B), suggesting that the KRAS binding peptides entered the cell, bound to KRAS, and disrupted the interactions with its signaling effectors. Control eGFP fusions whose membrane localization was not dependent on an interaction with KRAS (eGFP with a C-terminal farnesylation signal from HRAS, Fig S7A; eGFP with an N-terminal palmitoylation signal from Neuromodulin, Fig S7B) showed membrane accumulations with both binder and non-binder peptides.

**Figure 2:**
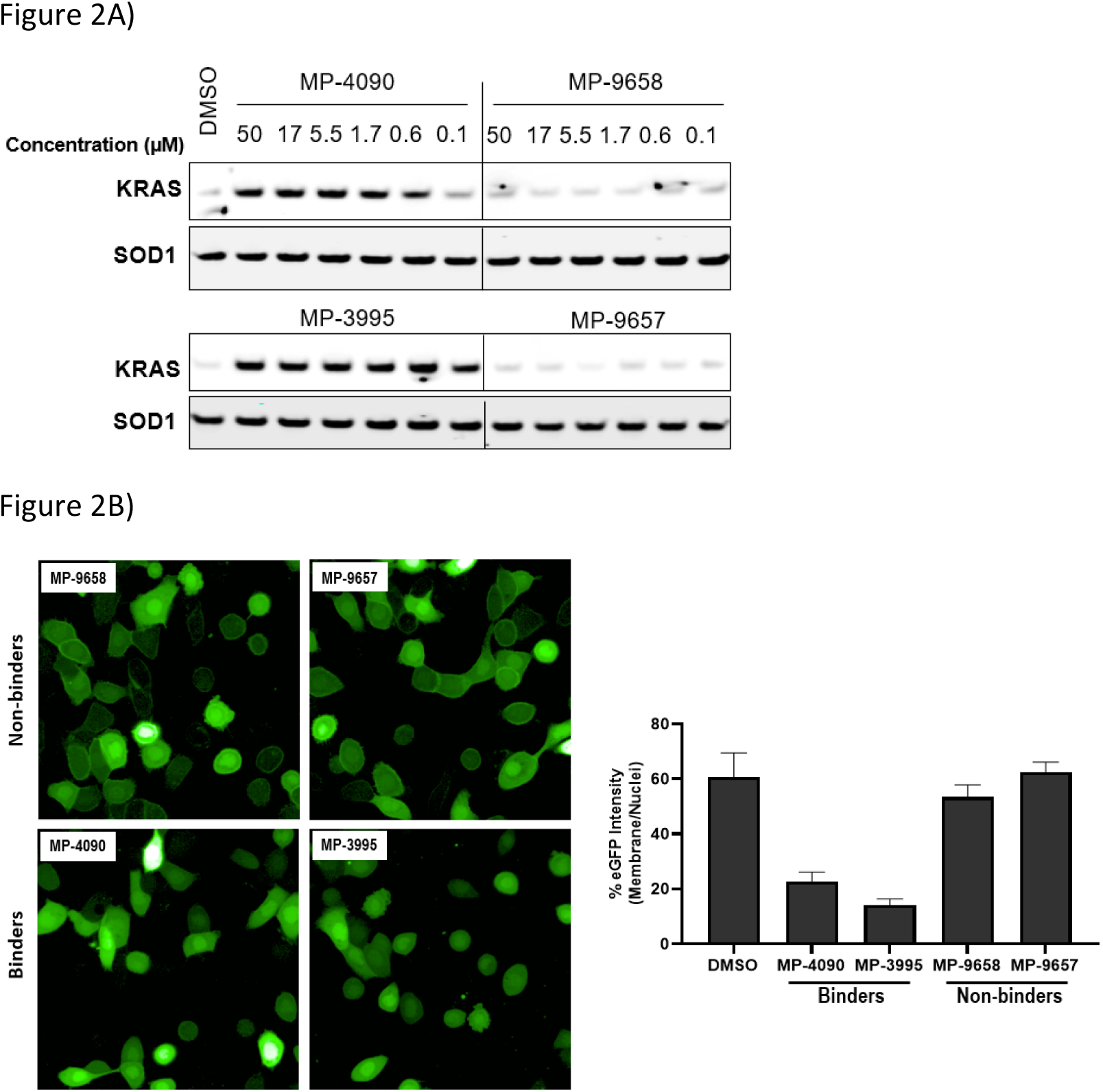
**MP-4090** and **MP-3995** specifically engages with cellular KRAS to inhibit its interactions with RAF–RBD. A) Isothermal CETSA experiments using intact AsPC-1 cells show that both **MP-4090** and **MP-3995** thermally stabilize KRAS^G12D^ whereas non-binding controls (**MP-9657** and **MP-9658**) do not. B) Introduction of a RBD-CRD-eGFP fusion protein into AsPC-1 cells (homozygous KRAS^G12D^) using mRNA transfection resulted in enrichment of the GFP signal at the membrane when cells were incubated with DMSO (not shown) or the non-binder control peptides, **MP-9657** and **MP-9658**. Membrane enrichment was lost when cells were incubated with **MP-4090** or **MP-3995**, suggesting that these peptides could effectively compete with the PPI.

### Exploring the mechanism of KRAS inhibition

We considered two distinct, non-mutually exclusive, mechanisms of action for this peptide series. First, the peptide might inhibit SOS-mediated nucleotide exchange, thus trapping KRAS in the inactive GDP-loaded state. Alternatively, although the peptide binding site does not overlap with the RAF RAS binding domain (RBD) binding site, the peptide might act as a GTP-state allosteric inhibitor to block the PPI and therefore KRAS signaling. Comparison of the KRAS–**MP-9903** complex with the KRAS–SOS (PDB 7KFZ) and KRAS–RBD (PDB 6XHB) complexes suggested that both mechanisms could potentially contribute to the inhibition of KRAS signaling. Specifically, binding of **MP-9903** should sterically/directly block the binding of SOS protein (Fig S8A). On the other hand, structural considerations suggest these peptides could act as allosteric inhibitors of RBD binding. Specifically, RBD binds at the PPI interface between the switch I and switch II regions, whereas **MP-9903** binds at the allosteric pocket between Switch II/helix H2 and helix H3. Upon binding of RBD, H2 moves towards H3 (Fig S8B), however binding of **MP-9903** moves the switch II and helix H2 towards the PPI interface. Such an outward conformation of switch II and helix H2 is not compatible for the binding of RBD, thus allosterically blocking RBD binding and KRAS signaling (Fig S8C, D). In agreement with this analysis and previous biochemical findings for **KRpep-2d** [10, 12, 13], our improved analogs blocked both GDP- and GTP-state activities. Specifically, **MP-3995** and **MP-9903** (one Arg on each terminus, Table 4) potently inhibited SOS mediated nucleotide exchange (Fig 3A). Compared to **KRpep-2d**, our improved analogs (**MP-6483, MP-4090, MP-3995**, and **MP-9903**) were also superior at inhibiting the interaction between KRAS^G12D^ and GST-RBD in a biochemical PPI assay (Fig 3B). We also probed PPI disruption with full-length, cellular KRAS, showing that **MP-6483** was highly effective at inhibiting the pulldown of endogenous b-RAF and c-RAF with HA-tagged KRAS^G12D^ from cell lysates (Fig 3C). Furthermore, studies with our alanine scan panel peptides (Table S1) demonstrated that PPI disruption efficiency corelated well with their TR-FRET binding affinities as effective blockade of RAF pull-downs was seen with binder peptides, with intermediate effects with the Leu7Ala analog, a peptide with moderate affinity. No effect was seen with non-binder peptides from the panel, the alanine mutants of Pro6, Ile9 or Asp12. The capacity to directly inhibit the GTP-loaded state was further supported by the lack of potency shift in our pERK assay when AsPC-1 or NCI-H358 (KRAS^G12C^) cells were treated with EGF (Fig S9A and S9B), a stimulus that shifts the nucleotide-loaded state towards GTP-bound KRAS. In contrast, the expected potency shift was observed with a GDP-state-selective G12C inhibitor (MRTX-1257) [3] in the presence of EGF treatment (Fig S9B). The capacity of these peptides to directly inhibit the GTP-loaded state was probed using HEK293 cells expressing either NanoLuc-KRAS^G12C^ or NanoLuc-KRAS^G12C/A59G^ under doxycycline control. A59G is a mutation that abrogates the remaining basal level of GTPase activity in KRAS^G12C^, thus pushing the protein more fully into the GTP state [20]. As expected, induced expression of the G12C and G12C/A59G mutant proteins led to increased KRAS signaling, as measured by pERK levels (Fig 3D). Treatment with the GDP-state preferring G12C inhibitor AMG 510 (sotorasib) resulted in covalent modification of the Nanoluc-KRAS^G12C^ target protein as evidenced by the increase in molecular weight on the Western blot (Fig 3D), something not seen with Nanoluc-KRAS^G12C/A59G^. AMG 510 treatment also blocked Nanoluc-KRAS^G12C^ but not Nanoluc-KRAS^G12C/A59G^ signaling, as indicated by pERK levels (Fig 3D). Together, these observations are consistent with the capacity of AMG 510 to exclusively bind to and inhibit GDP-loaded KRAS^G12C^ and for the A59G mutation to further bias the protein to the GTP state. In contrast, **MP-3995** was effective at inhibiting both the single and double mutants (Fig 3D), suggesting that the peptide can inhibit signaling even when KRAS is pushed more completely into the active conformation.

**Figure 3:**
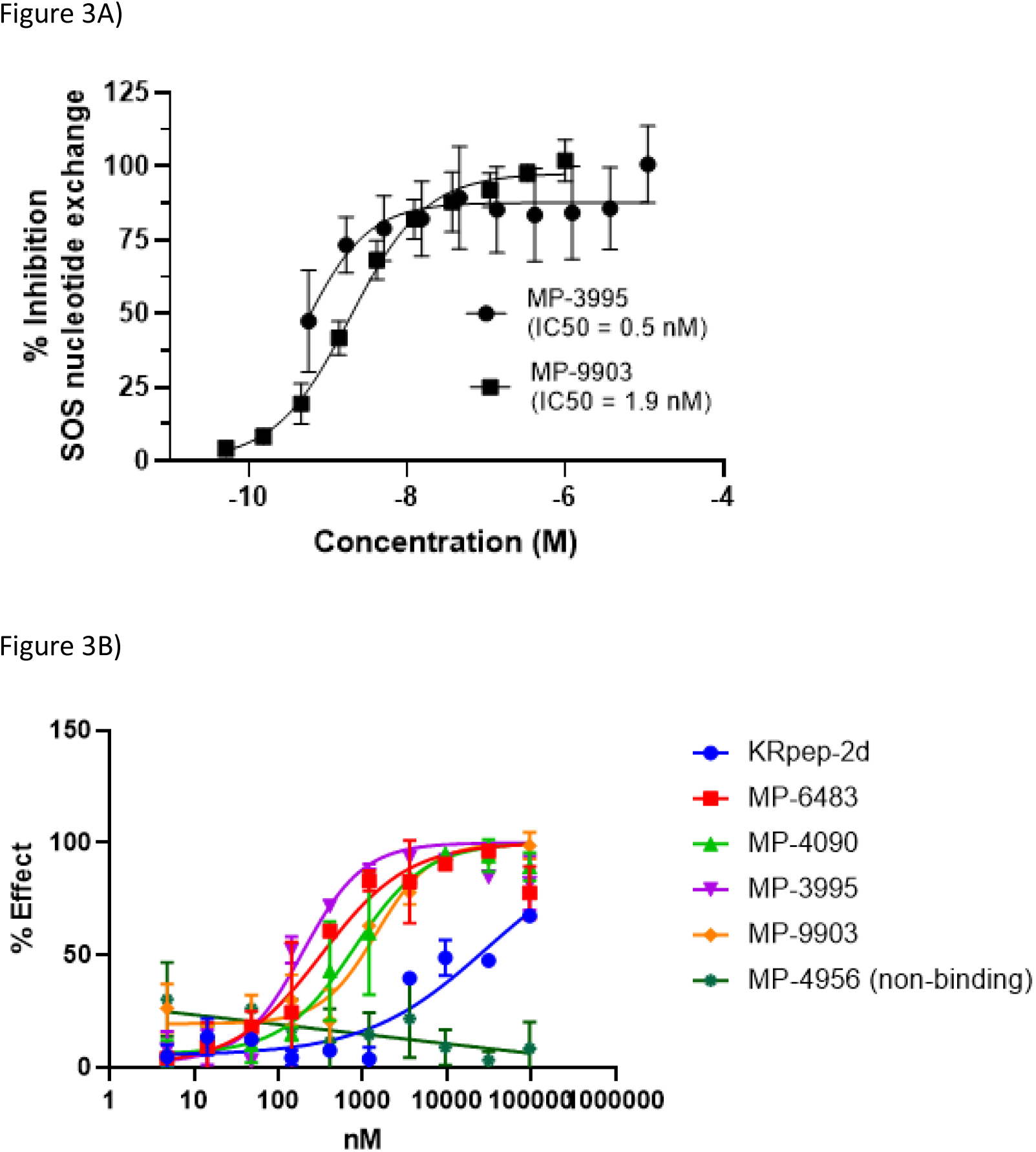
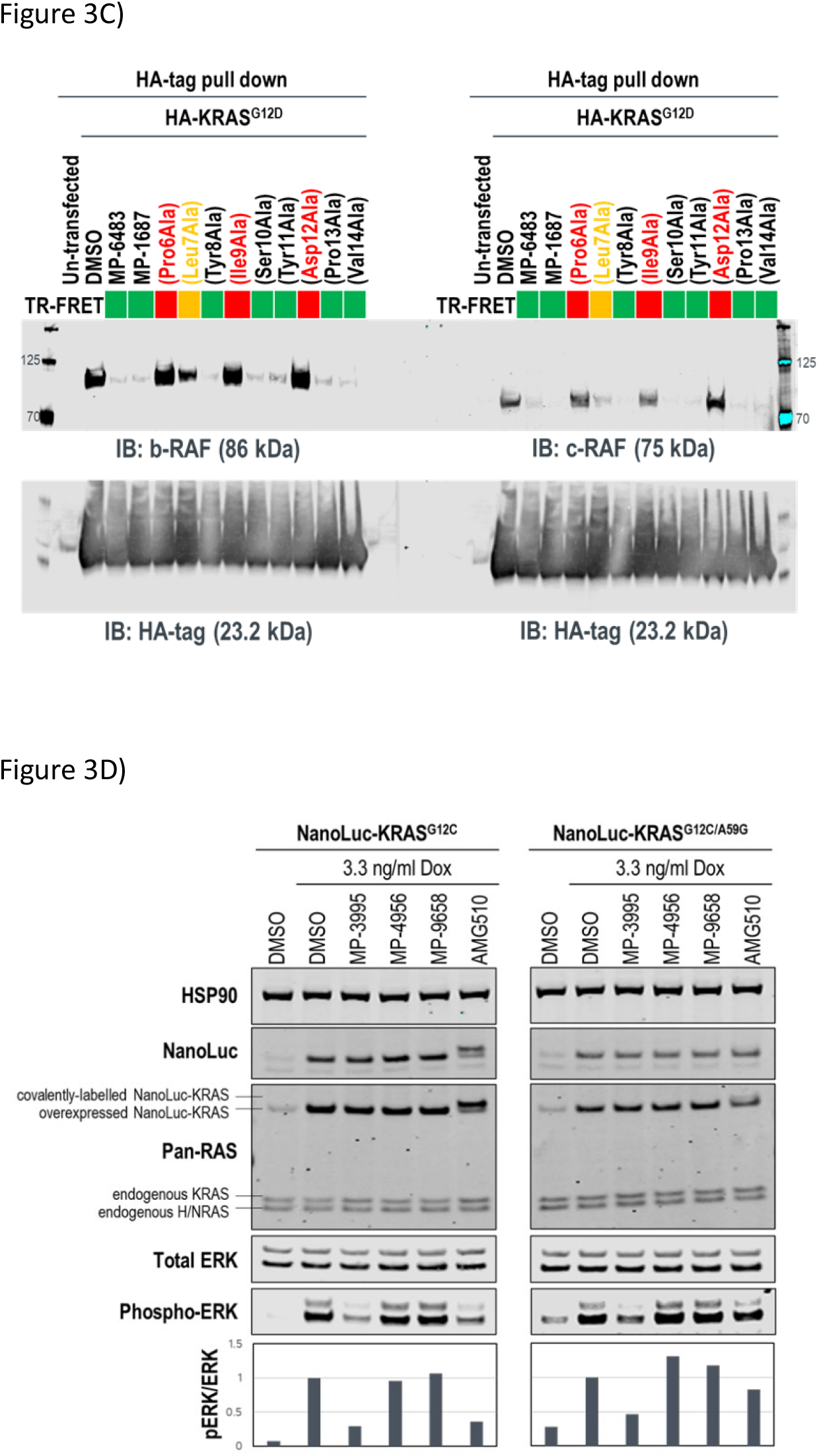
**KRpep-2d** peptide analogs have dual inhibitory mechanisms. A) **MP-3995** and **MP-9903** potently inhibited SOS-mediated nucleotide exchange. B) **MP-6483, MP-4090, MP-3995**, and **MP-9903** showed a superior capacity to block the KRAS/RBD PPI compared to **KRpep-2d**, whereas the non-binder control (**MP-4956**) had no activity. C) **MP-6483** and **MP-1687** blocked the co-immunoprecipitation of b-RAF (left panel) and c-RAF (right panel) with KRAS^G12D^. Alanine mutant peptide library analogs of **MP-1687** (Table S1) demonstrated that KRAS binding affinity correlated well with disruption of the PPI. D) **MP-3995** blocked phospho-ERK signaling in cells expressing either NanoLuc-KRAS^G12C^ or NanoLuc-KRAS^G12C/A59G^ whereas AMG510 only inhibited NanoLuc-KRAS^G12C^, non-binders (**MP-4956** and **MP-9658**) had no activity.

### MP-3995 inhibits pERK and cell proliferation in a panel of mutant KRAS cell lines

Next, we probed whether the dual inhibitory mechanism of **MP-3995** could translate into pERK inhibition across a panel of cells lines. In a variety of KRAS^G12C^, KRAS^G12V^, and KRAS^G12D^ cancer lines, **MP-3995** blocked pERK signaling in the low micromolar range (Fig 4A), thus demonstrating this molecule is a pan-KRAS inhibitor. These effects also translated to inhibitory effects on cell proliferation in a KRAS^G12D^ line (AsPC-1 cells) as well as lines harboring KRAS^G12V^ (SK-CO-1) and KRAS^G12C^ (NCI-H358 and NCI-2122) (Fig 4B). Importantly, the all-D non-binding control peptide, **MP-4956**, had minimal inhibitory effects in these lines, suggesting an on-target effect (Fig 4B). Mostly importantly, **MP-3995** had no effect on pERK inhibition in A375 cells (Fig 4A), consistent with the lack of cell growth inhibition by the peptide in this line and the on-target nature of the molecule (Fig 4B). In a separate experiment, we identified different cell lines sensitive and insensitive to **MP-3995** (Fig 4C). For the latter, this included HEK293 cells (Fig 4C), a cell line that has no dependency on RAS for cell proliferation [21]. Overall, this extended cell proliferation panel showed cell proliferation effects for eight out of thirteen cell lines.

**Figure 4:**
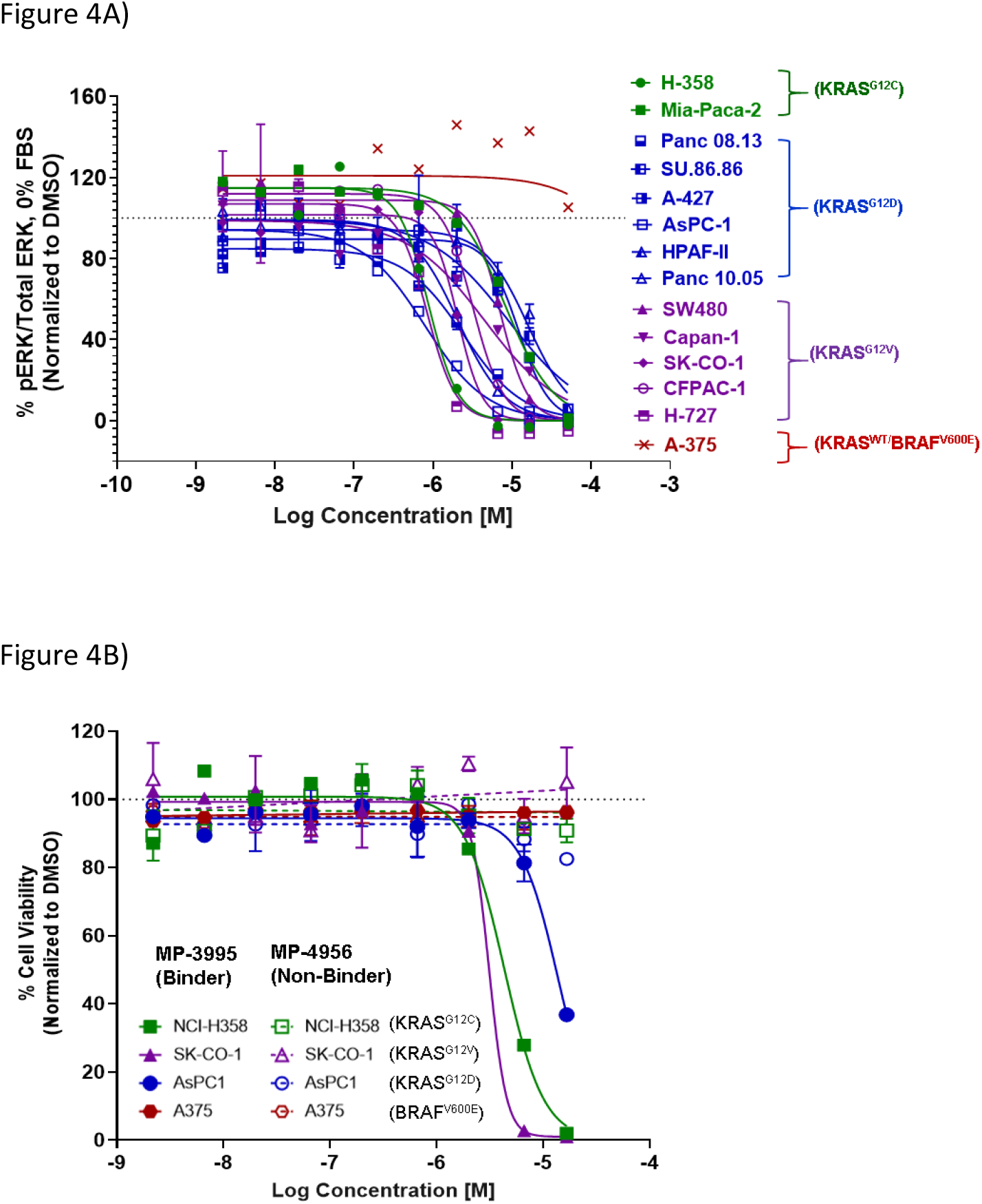
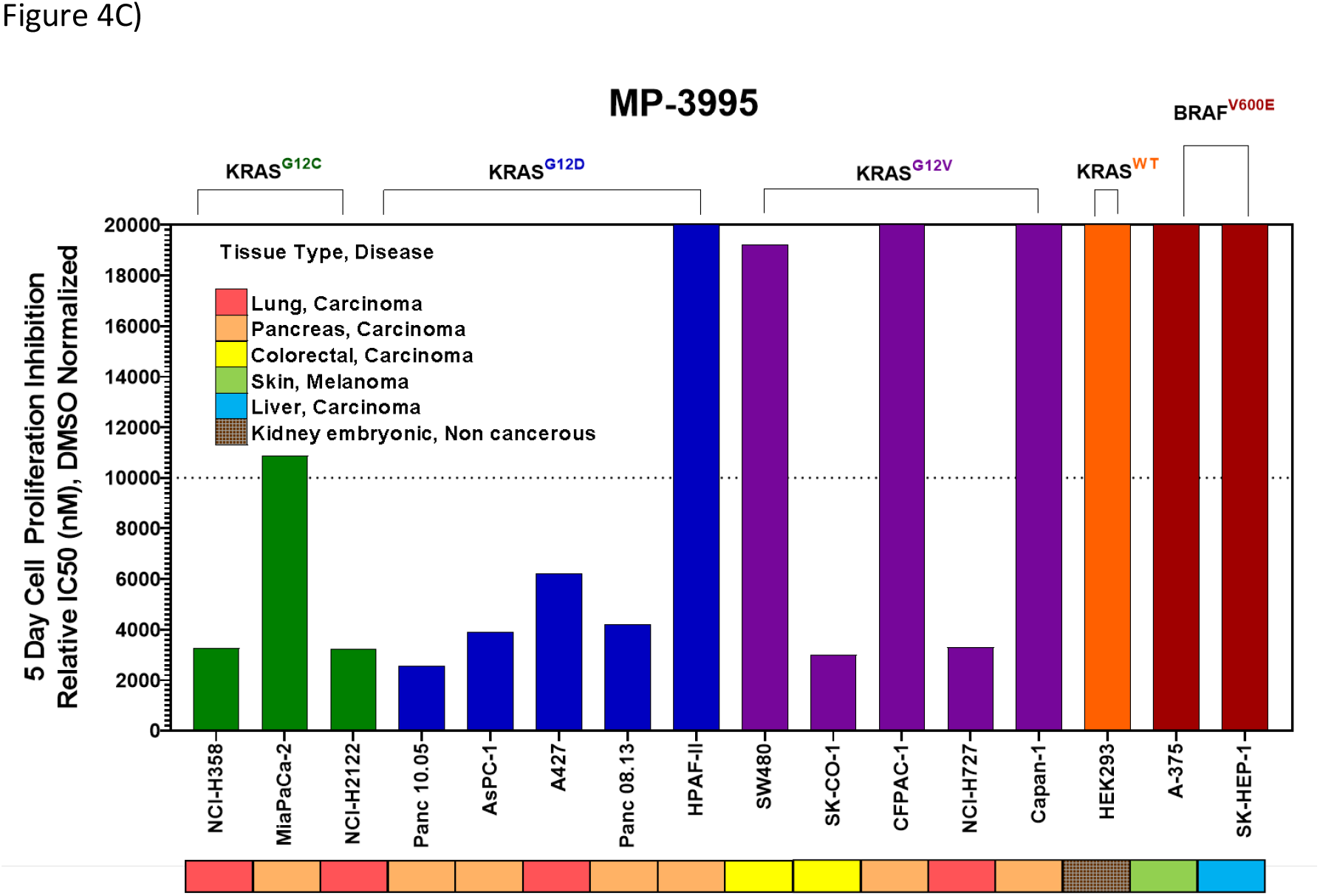
**MP-3995** inhibits cell proliferation in cell lines of different tissue origins bearing various KRAS mutations. A) **MP-3995** inhibits phospho-ERK in KRAS mutant cell lines but not BRAF^V600E^ (KRAS independent) A375 cells. B) **MP-3995**, but not the non-binder control **MP-4956**, inhibits cell proliferation in KRAS^G12D^, KRAS^G12C^ and KRAS^G12V^ mutant cells. C) **MP-3995** inhibits cell proliferation in 8 out of 13 KRAS mutant cell lines across different disease indications.

### The induction of mast cell degranulation by arginine-rich KRpep-2d analogues represents a barrier to their progression as therapeutics

Cationic peptides can induce IgE-independent mast cell degranulation (MCD, [22]), a process involving the release of biogenic amines (most notably histamine), as well as a cocktail of proteases, cytokines, leukotrienes and prostaglandins [23]. This can result in itchiness, hives, edema, and even death through anaphylactic shock. Unfortunately, **KRpep-2d** (data not shown) and analogues, including **MP-4090**, proved to be highly potent activators of MCD, as determined by an *ex vivo* histamine release assay using rat mast cells. To investigate further, we probed histamine release for a series of **MP-4090** peptides containing varying numbers of arginines on the N- and C-termini (Table 4). The reported rat mast cell degranulation (rMCD) value in Table 4 is a threshold for the lowest concentration at which a two-fold change in histamine release was observed. Arginine content correlated with the level of histamine release in these cells, with peptides containing 0–2 arginines (**MP-9903, 15** and **16**) being free from this potential liability. Interestingly, MCD activity was observed to be inversely correlated to peptide permeability as monitored by the NanoClick assay, leading to a lack of cellular activity when the Arg-count was less than 8. The MCD liability is independent of the stereochemistry of the arginine tails. Of note, the binding affinity was minimally affected by the number and stereochemistry of arginine residues.

## Discussion

The recent landmark approval of sotorasib (LUMAKRAS™) for KRAS^G12C^ mutated non-small cell lung cancer (NSCLC), provides definitive pharmacological validation of this target for human cancers. This long-sought victory will undoubtedly benefit a subset of patients. However, for the large number of cancers driven through non-G12C mutant KRAS (e.g. G12D and G12V), much work remains. For these mutations, leveraging the switch II pocket with non-covalent analogs is an obvious approach. However, it is uncertain whether this strategy will be successful. As well, given the importance of KRAS and its associated challenges, orthogonal approaches need to be pursued.

The discovery of **KRpep-2d**, a macrocyclic peptide with validated high-affinity binding, represented an attractive starting point for the identification of a non-G12C KRAS inhibitor. However, we recognized that the disulfide-mediated macrocyclization strategy represented a barrier to cellular activity since it would likely be rapidly reduced leading to the non-binding linear form in the intracellular compartment. Indeed, in our hands, this molecule had no effect on cellular KRAS signaling and exhibited a very short half-life in cell homogenate. Most attempts to replace the disulfide bond with a non-reducible linkage resulted in peptides with forfeited KRAS binding. However, by extending our search to include variation of stereochemical centers, we discovered that the linkage of a D-Cys5 residue to Cys15 through a thioacetal bridge resulted in a redox stable, high affinity binder. Next, replacing the proteolytically unstable N- and C-terminal tetra-arginine tails with their enantiomeric counterparts, resulted in a peptide with cellular activity. Addressing a metabolic soft spot within the macrocycle through a S10 to α-methyl-Ser substitution further improved metabolic stability and consequently sustained cellular activity. Such constrained building blocks can force peptides into their biologically active conformations, while often providing remarkable resistance to enzymatic degradation.

Studies by us and others [10-13] suggest that the **KRpep-2d** peptides series might inhibit KRAS signaling in at least two distinct ways, by directly blocking the interaction with KRAS effectors (e.g., RAF) as well as by indirectly preventing these interactions by blocking the conversion of the GDP (off) state to the GTP (on) state. Both activities may indeed be contributing to cellular KRAS inhibition. Dual inhibition of mutant KRAS signalling is attractive, especially considering the observation that cancer cells can reactivate the MAPK pathway to resist G12C covalent inhibitors [24], molecules that trap the protein in the GDP state [21]. Initial studies suggest that such compensatory mechanisms may not occur with **KRpep-2d** family members. First, pathway up-regulation with EGF stimulation did not alter the potency of **MP-4090** (Fig S9). In addition, biasing KRAS protein to the GTP (on) state with the G12C/A59G double mutant prevented pERK inhibition with AMG 510 (sotorasib) but not **MP-3995** (Fig 3D), suggesting that this peptide can continue to prevent mutant KRAS signaling even when the protein is pushed into the active state.

A key foundation for the identification of *bona fide*, functionally active peptides targeting intracellular proteins is the application of stringent experimental controls, both chemical and biological in nature. This is important as cationic and hydrophobic elements that are often highly prevalent in cell penetrant peptides can lead to false positive cellular read-outs through cell membrane disruption. Indeed, this has been shown specifically for peptides that have been incorrectly reported to have on-target cellular activity against KRAS [15]. Recently, it was demonstrated that these features can also lead to phospholipidosis, at least when present in small molecules [25]. Thus, we applied a host of chemical and biological controls in our studies. For the former, we made a series of control peptides designed to be as similar as possible to the cell active molecules but devoid of KRAS binding. This was accomplished through stereo-inversion of all or key binding amino acids. The observation that these peptides were indeed non-binders but also inactive in our pERK assay suggested that the cell activity of our lead peptides (**MP-4090** and **MP-3995**) was due to the consequences of KRAS binding. A series of biological controls gave us further confidence that these peptides had *bona fide* cellular activity. Indeed, the absence of LDH release and off-target activity in an irrelevant signaling pathway was consistent with the on-target nature of this series. As well, the lack of pERK and proliferation inhibition activities in a KRAS independent line (A375) further suggested that off-target mechanisms were not involved. In addition, evidence of direct target engagement was provided by peptide-induced KRAS thermal stabilization in a CETSA assay and through the capacity of these peptides to displace GFP–RBD–CRD protein from the cell membrane. In both cases, non-binding control peptides gave negative results, as expected.

The reliance on poly-arginine sequences at the N- and C-termini make it challenging to progress these peptides towards the clinic. Indeed, *ex vivo* mast cell degranulation studies confirmed a strong correlation between the number of arginine residues and histamine release, a phenomenon that can lead to serious consequences *in vivo*. Unfortunately, reducing the number of arginines also correlated with a loss of cell permeability and subsequent cellular activity, therefore confounding their pipeline progression.

Four decades of KRAS research has started to bear fruit with the 2021 approval of the first direct inhibitor of a specific KRAS mutation, G12C. The search for efficacious inhibitors of all other mutant KRAS-driven cancers continues. Despite the structural liabilities identified by us, the peptides described herein represent valuable templates for achieving in vivo activity against KRAS-driven, non-G12C cancers. These efforts would focus on identification of sequence variants that achieve cell entry without dependence on arginine-rich sequences. Our efforts directed towards such objectives will be the subject of future communications.

Overall, systematic studies reported here made key advances on this peptide series and used rigorous controls to validate the potential of blocking mutant-KRAS function in cancer cells via binding to this unique epitope. As such, novel avenues are open for impacting KRAS-mediated cancers beyond the recent successes with covalent G12C modulation.

## Acknowledgements

We thank Villa Zheng, Mike Xue, and Kyle Chen at Chinese Peptide Company (CPC) for peptide synthesis support. We thank Dr. Jennifer Johnston for help with figure preparation.

## TOC Graphic

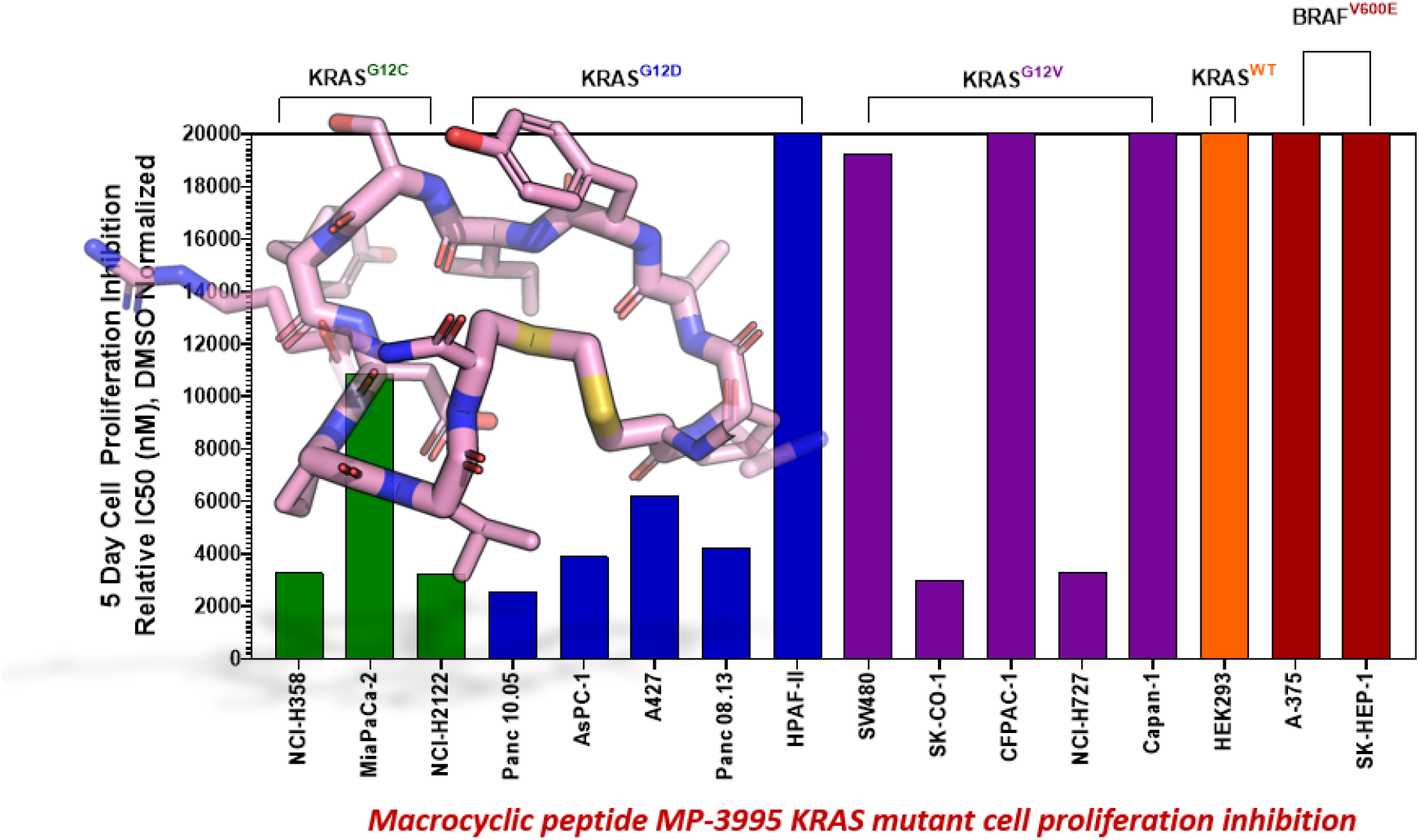

### Synopsis

Novel on-target macrocyclic peptide inhibitors of KRAS show broad inhibition of proliferation of multiple KRAS-dependent cancer cell lines beyond recently clinically approved G12C inhibitor.

