## Supplemental Information for "Discovery of cell active macrocyclic peptides with on-target inhibition of KRAS signaling"

Shuhui Lim[*, 1], Nicolas Boyer[*, 2], Nicole Boo[1], Chunhui Huang[2], Gireedhar Venkatachalam[1], Yu-Chi Angela Juang[1], Michael Garrigou[2], Kristal Kaan[1], Ruchia Duggal[2], Khong Ming Peh[1], Ahmad Sadruddin[1], Pooja Gopal[1], Tsz Ying Yuen[3], Simon Ng[3], Srinivasaraghavan Kannan[3], Christopher J. Brown[3], Chandra Verma[3], Peter Orth[4], Andrea Peier[4], Lan Ge[4], Xiang Yu[5], Bhavana Bhatt[5], Feifei Chen[5], Erjia Wang[5], Nianyu Jason Li[5], Raymond J. Gonzales[5], Alexander Stoeck[2], Brian Henry[1], Tomi Sawyer[2], David Lane[3], Charles W. Johannes[ψ, 3], Kaustav Biswas[ψ, 2], Anthony W. Partridge[ψ, 1]

[1] MSD International, Singapore 138665, Singapore.

[2] Merck & Co., Inc., Boston, Massachusetts 02115, United States.

[3] Agency for Science, Technology and Research (A*STAR) Singapore 138665, Singapore.

[4] Merck & Co., Inc., Kenilworth, New Jersey 07033, United States.

[5] Merck & Co., Inc., West Point, Pennsylvania, 19486, United States

[*] These authors contributed equally to this work

**Supplemental Information table of contents**


- Supplementary figures S1 – S9 ……………………………………………………………………………………………pages S3 – S14
- Supplementary Tables S1 – S4 ……………………..……………………………………………..……………………………...pages S15 – S19
- Materials and methods ……………………………………………………………………………..…………………………….pages S20 – S41
- Supplemental references ………………………………………………..…………………………………………………..page S42

**Supplementary Figures:**

S1)

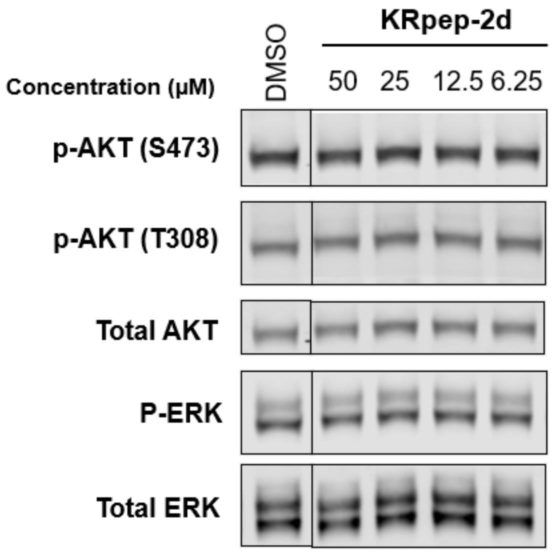


**Figure S1: KRpep-2d** does not show inhibition of phospho-ERK signaling in AsPC-1 (KRAS$^{G12D}$) cells, as assessed by Western blot. Lane 1 is DMSO control.

S2)

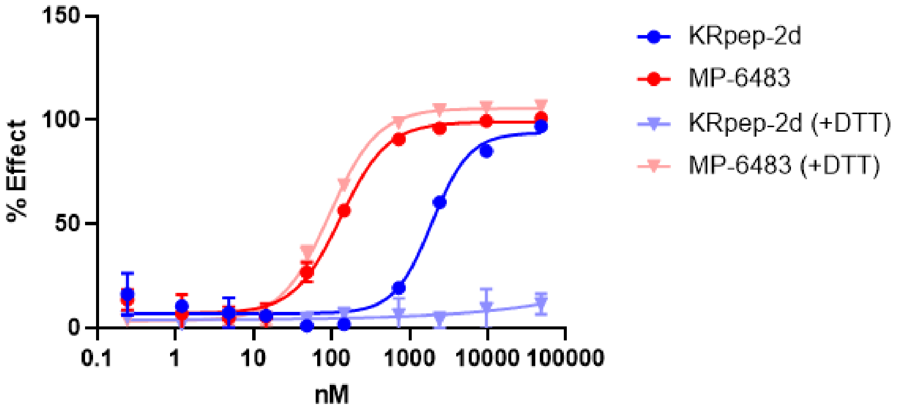


**Figure S2: KRpep-2d** (redox-sensitive macrocycle) loses binding to GDP-loaded KRAS$^{G12D}$ in the presence of DTT, whereas **MP-6483** (redox-stable macrocycle) maintains binding, as assessed by TR-FRET assay.

S3)

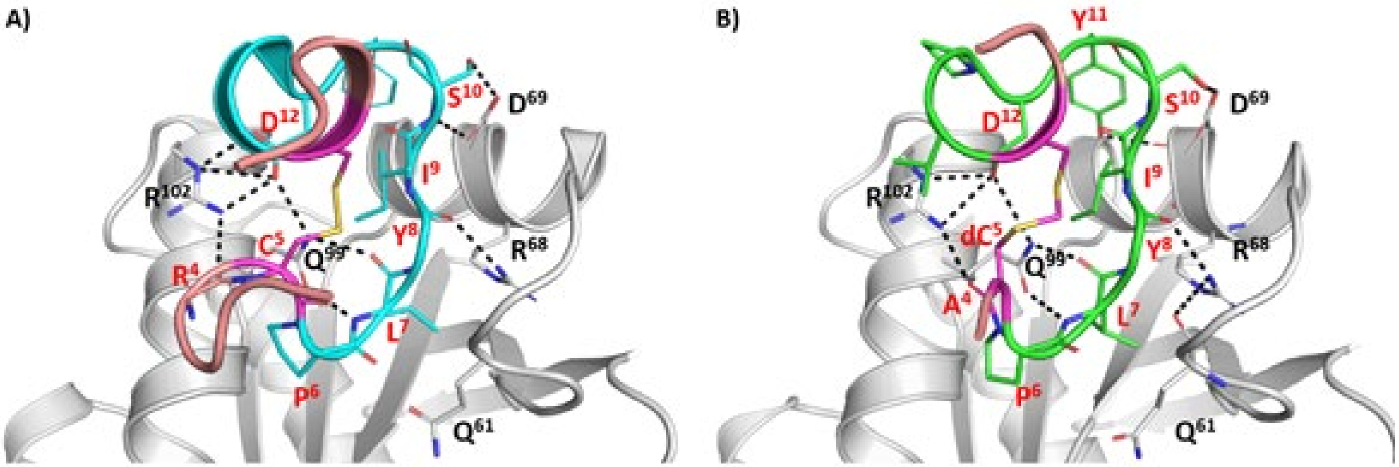


**Figure S3:** Structures of KRAS – peptide complexes, crystal structure of GDP-loaded KRAS$^{G12D}$ in complex with **KRpep-2d** (A) or molecular dynamics structure of GMPPCP-loaded KRAS$^{G12D}$ in complex with **MP-9903** (B). KRAS is shown in grey coloured ribbon and the bound peptide is shown as cartoon (Cyan for **KRpep-2d** and Green for **MP-9903**) with the terminal Arg (Salmon), methylene bridge (magenta) and interacting residues highlighted as sticks. H-bond interactions are highlighted as dashed lines (black).

S4)

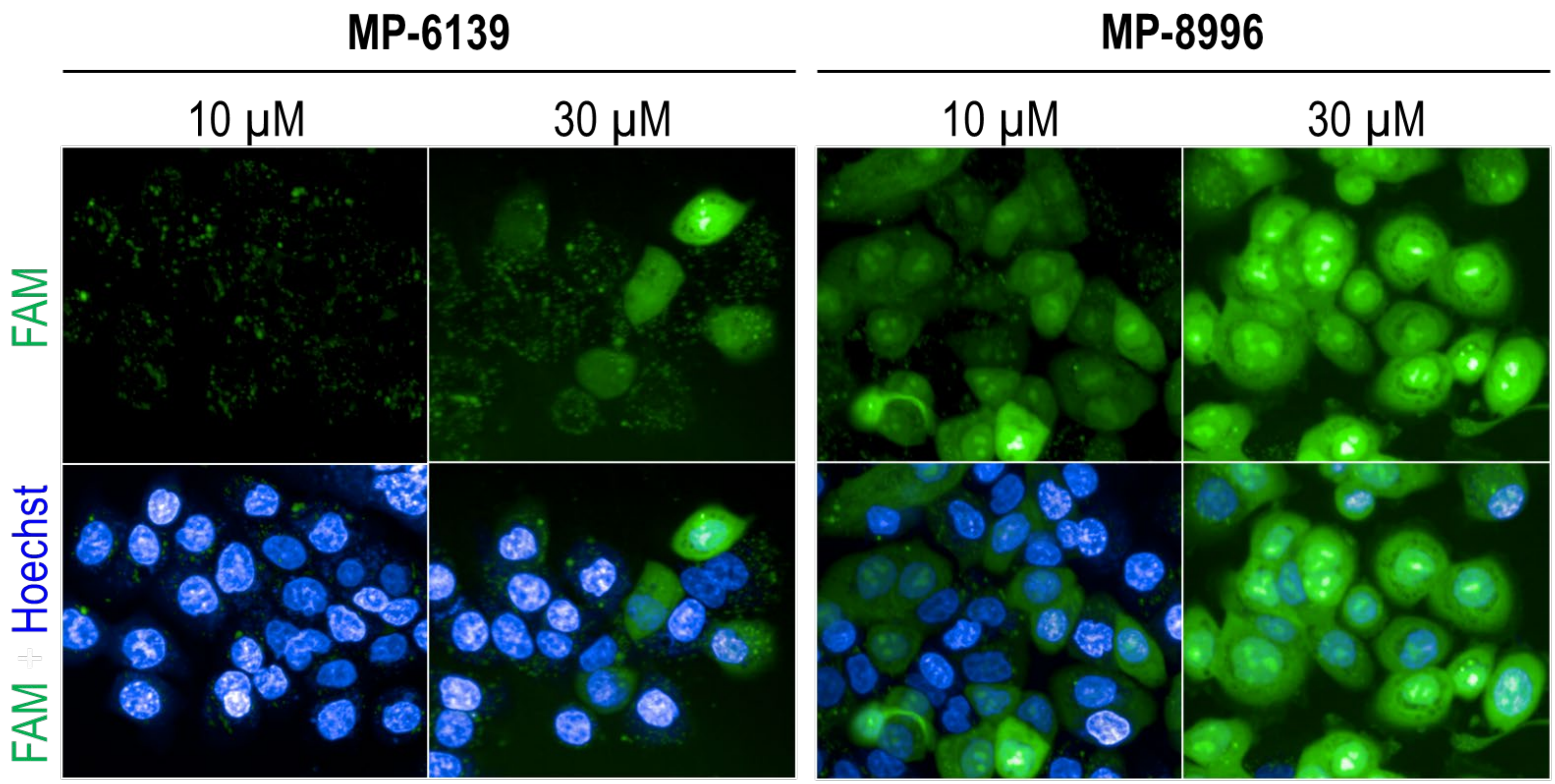


**Figure S4:** Substitution of L-arginines (**MP-6139**) for D-arginines (**MP-8996**) resulted in a marked increase in uptake in AsPC-1 cells (30 min peptide treatment), as assessed by confocal imaging using FAM-labeled peptides. See Table S4 for full sequences of the FAM-labeled analogs.

S5A)

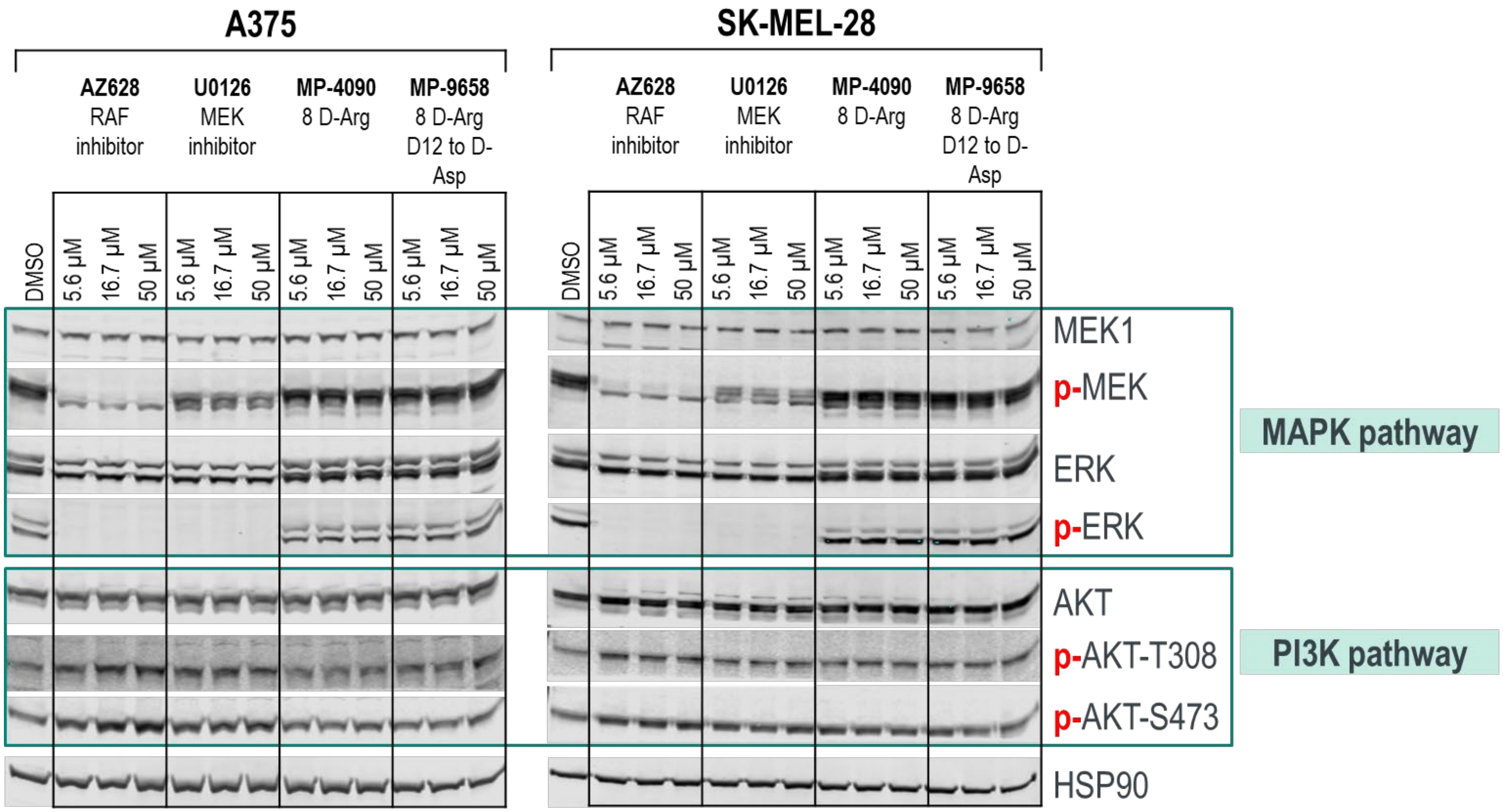
A375
SK-MEL-28
AZ628
RAF
inhibitor
U0126
MEK
inhibitor
MP-4090
8 D-Arg
MP-9658
8 D-Arg
D12 to D-Asp
DMSO
5.6 µM
16.7 µM
50 µM
MEK1
p-MEK
ERK
p-ERK
AKT
p-AKT-T308
p-AKT-S473
HSP90
MAPK pathway
PI3K pathway

S5B)

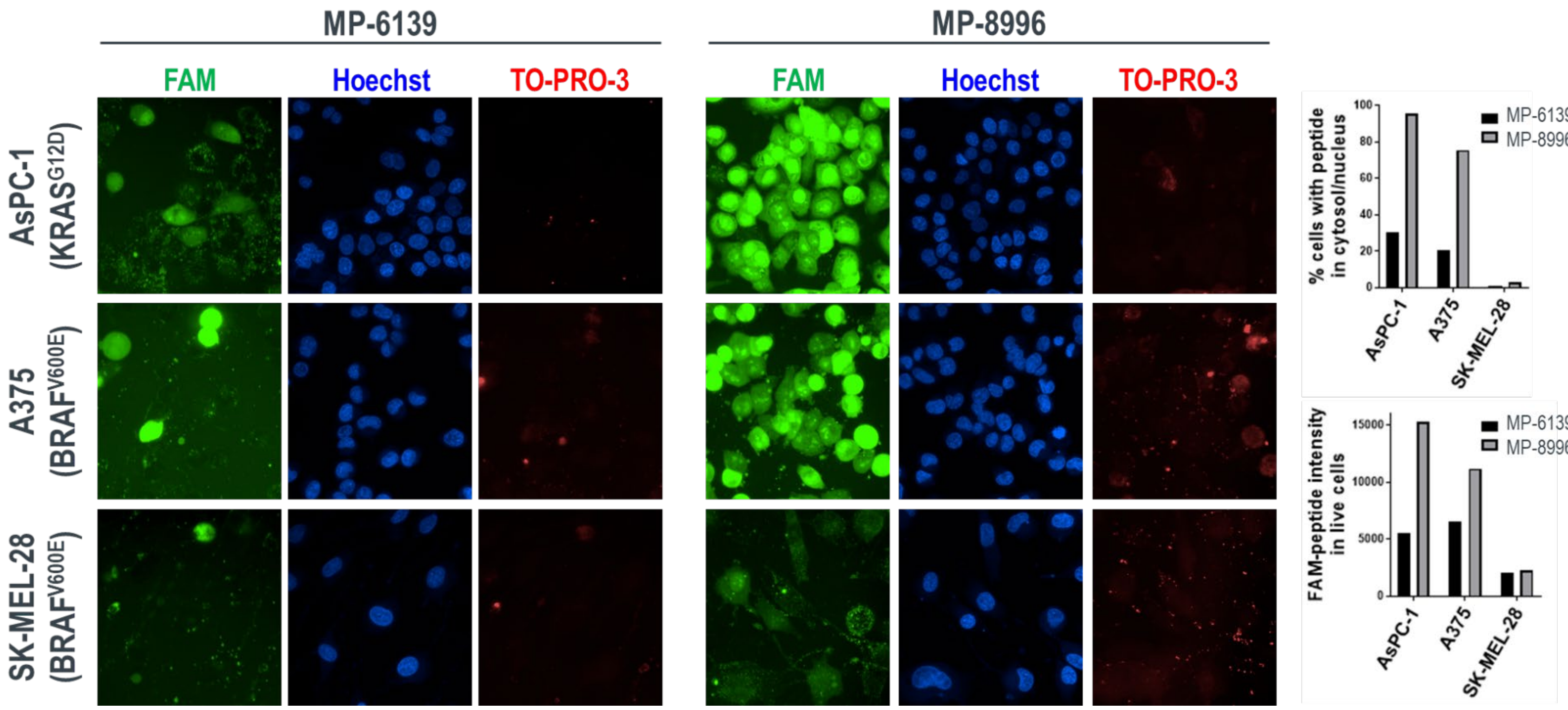


**Figure S5:** Peptides with D-arginines have increased cell permeability but do not inhibit MAPK and PI3K signaling in cancer cell lines with BRAF mutations. A) **MP-4090** does not inhibit MAPK and PI3K signaling in A375 and SK-MEL-28 cell lines. B) A FAM-labeled version of **MP-4090** (**MP-8996**) entered these cells with roughly equivalent amounts compared to AsPC-1 cells. **MP-6139** is the FAM-labeled version of the corresponding L-Arg peptide **MP-6483**.

S6A)

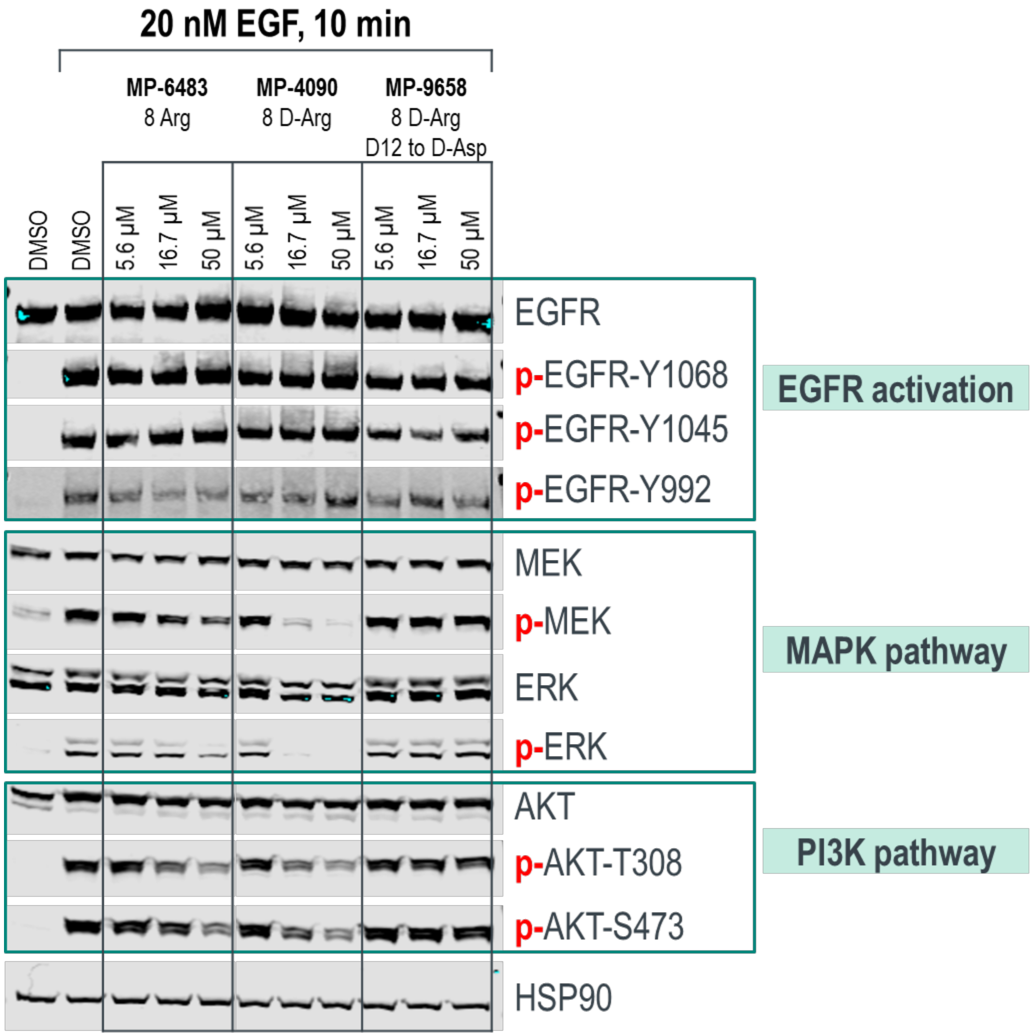

20 nM EGF, 10 min
MP-6483
8 Arg
MP-4090
8 D-Arg
MP-9658
8 D-Arg
D12 to D-Asp
DMSO
DMSO
5.6 µM
16.7 µM
50 µM
5.6 µM
16.7 µM
50 µM
5.6 µM
16.7 µM
50 µM
EGFR
p-EGFR-Y1068
p-EGFR-Y1045
p-EGFR-Y992
EGFR activation
MEK
p-MEK
ERK
p-ERK
MAPK pathway
AKT
p-AKT-T308
p-AKT-S473
PI3K pathway
HSP90

S6B)

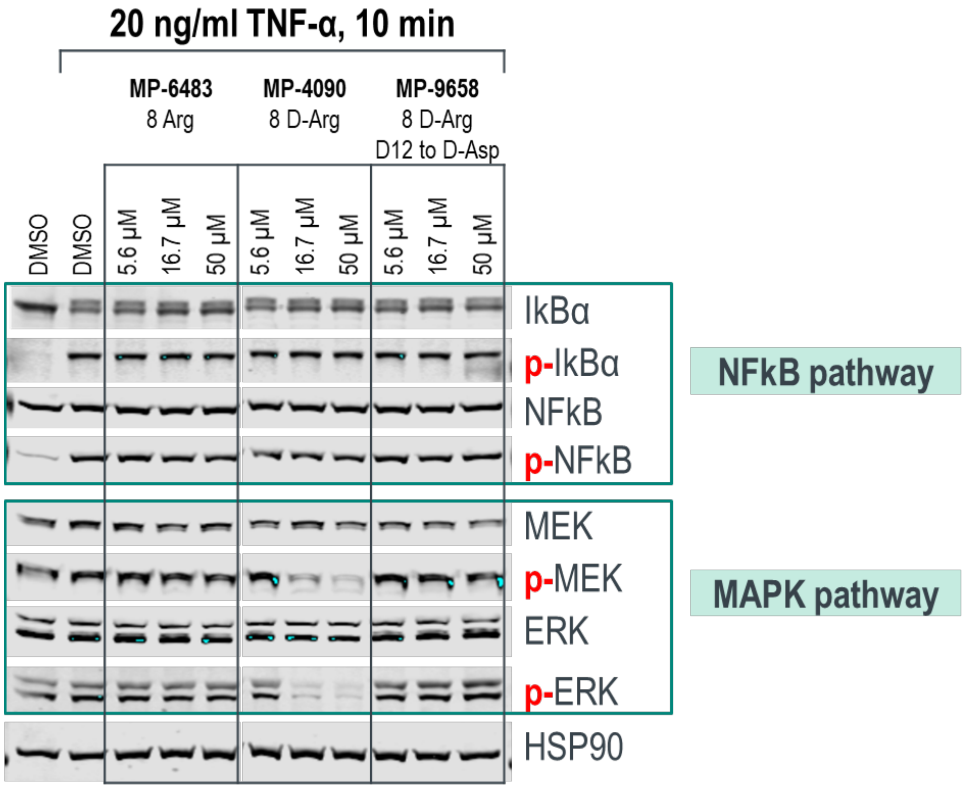


**Figure S6:** Inhibition by redox-stable KRpep-2d family members is specific to signaling events downstream of KRAS. A) **MP-6483** and **MP-4090** inhibit signaling downstream of KRAS (pERK, pMEK, and pAKT) but not upstream of KRAS (pEGFR). B) **MP-6483**, **MP-4090**, and non-binding control **MP-9658** have no effect on NFκB signaling.

S7A)

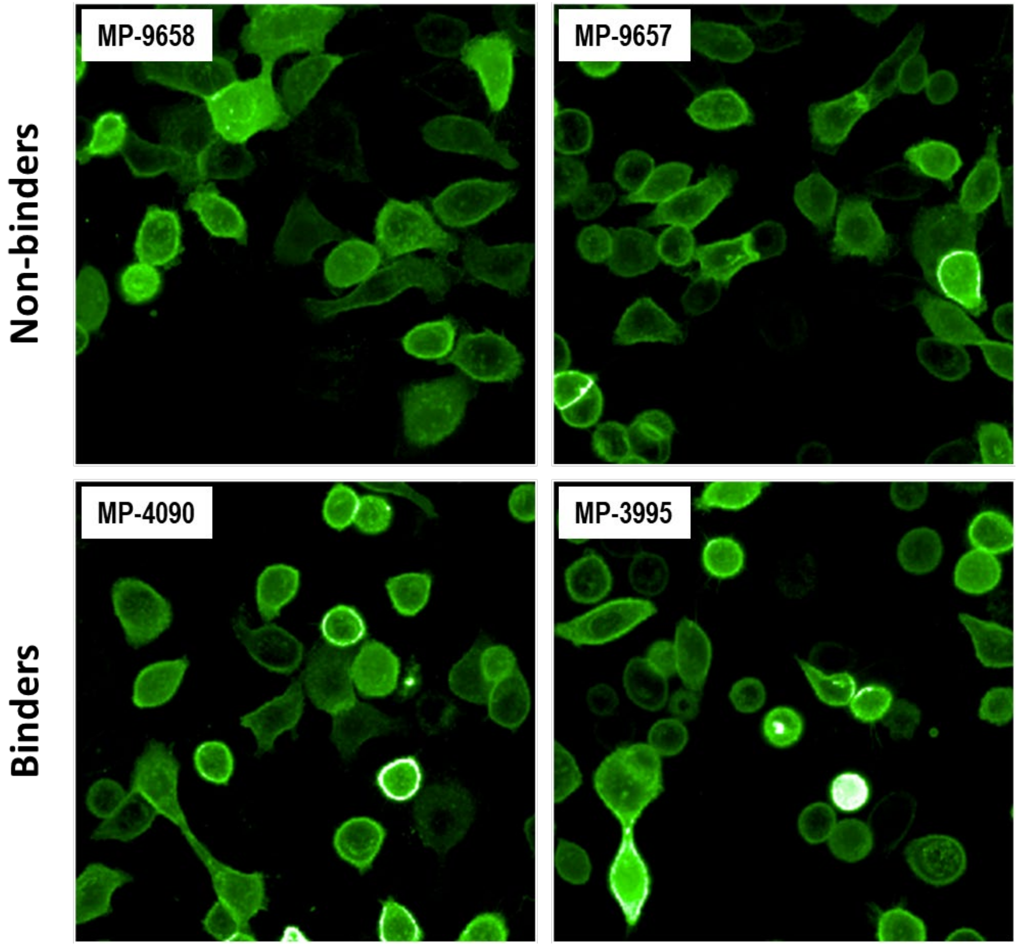

S7B)

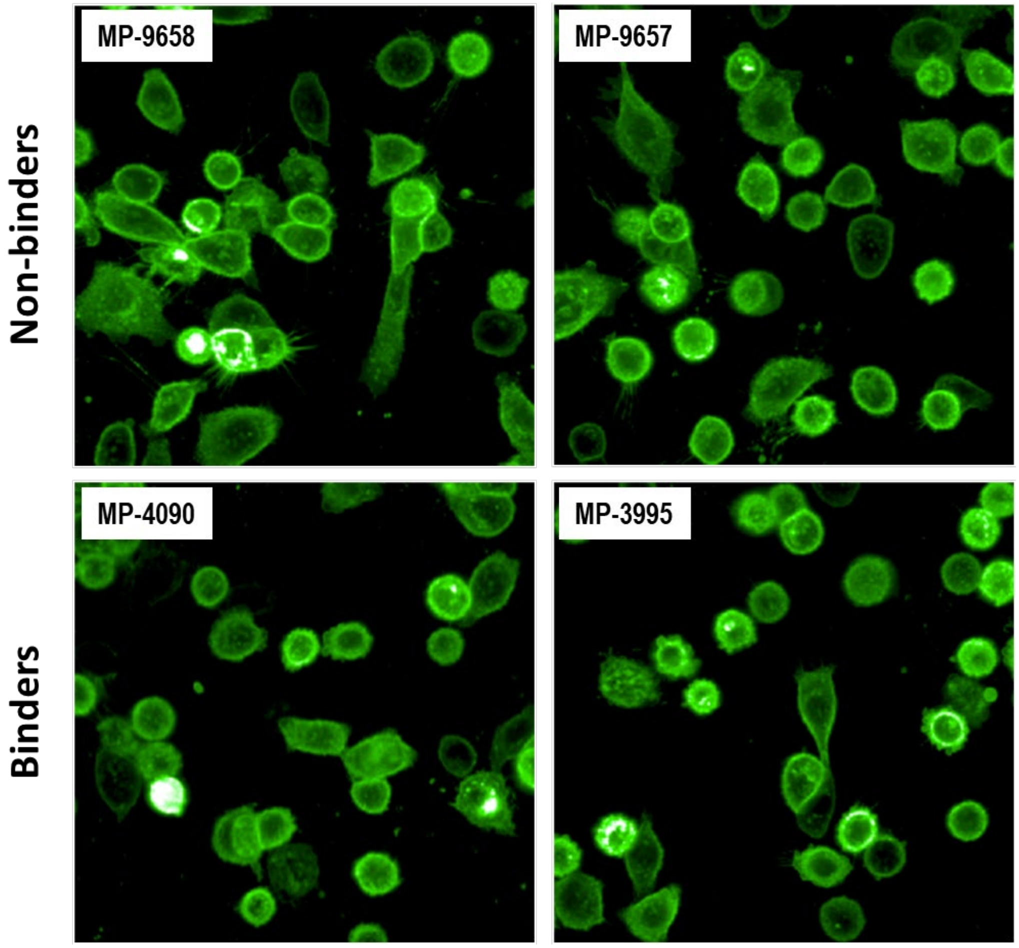


**Figure S7:** eGFP fusions whose membrane localization is not dependent on KRAS binding are not displaced from the membrane by **MP-4090**, **MP-3995**, nor by non-binding controls. A) Cells expressing eGFP fused at the C-terminal to the farnesylation signal from HRAS. B) Cells expressing a protein consisting of the N-terminal palmitoylation signal from Neuromodulin fused to the N-terminus of eGFP.

S8)

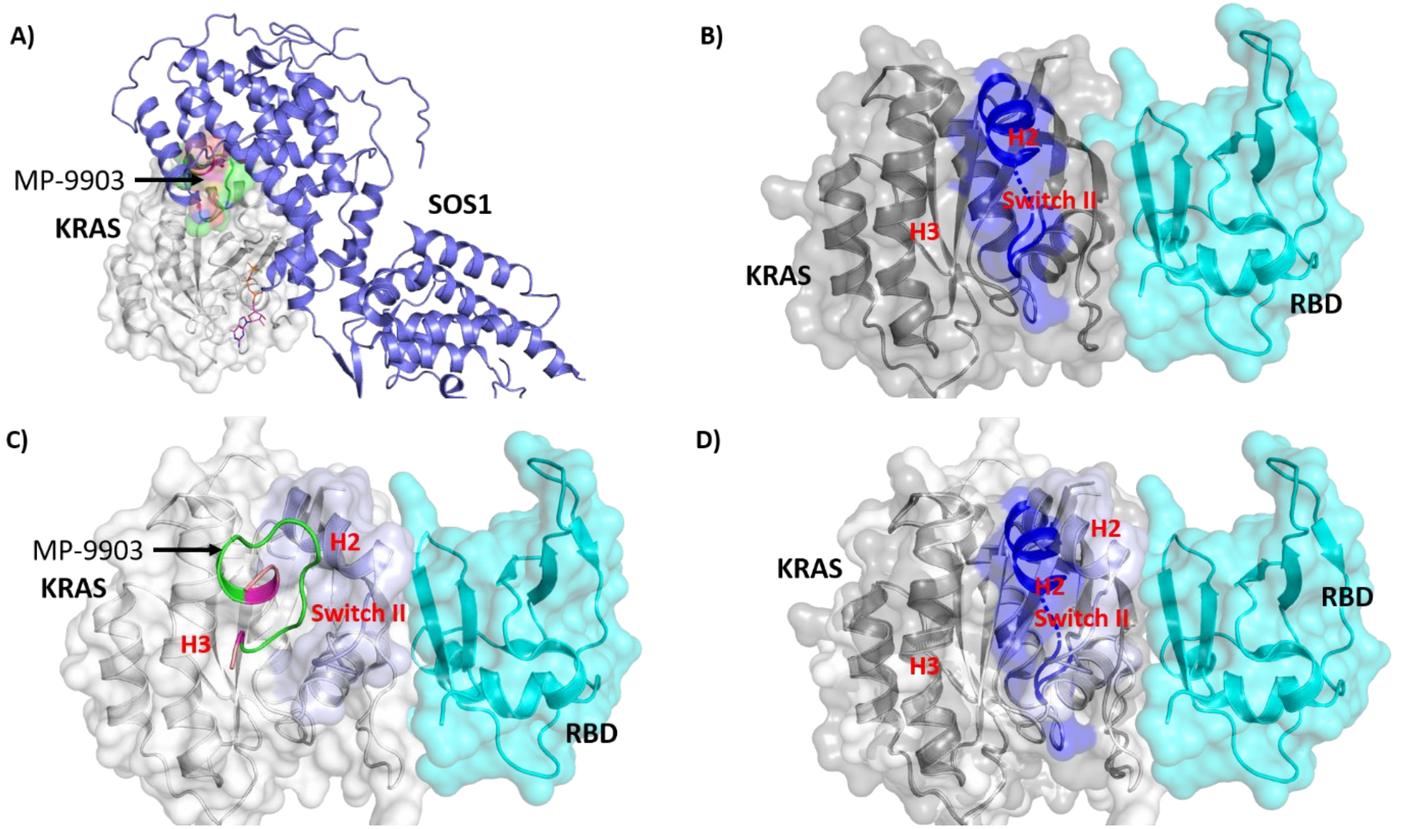


**Figure S8:** Comparison of the crystal structure of KRAS–**MP-9903** with the crystal structure of KRAS–SOS/RBD complexes (PDB ID's 7KFZ and 6XHB). (A) KRAS is shown in grey cartoon/surface, **MP-9903** is shown in green cartoon/surface and SOS1 is shown in blue cartoon. (B) KRAS is shown in dark grey cartoon/surface and RBD is shown in teal cartoon/surface, highlighting the inward movement of helix H2 and switch II (C) KRAS is shown in grey cartoon/surface, **MP-9903** is shown in green cartoon, highlighting the outward movement of helix H2, and switch II, induced by the binding of **MP-9903** (D) overlay of KRAS–**MP-9903** (peptide not shown) and the KRAS–RBD complex (colouring scheme same as B and C).

S9A)

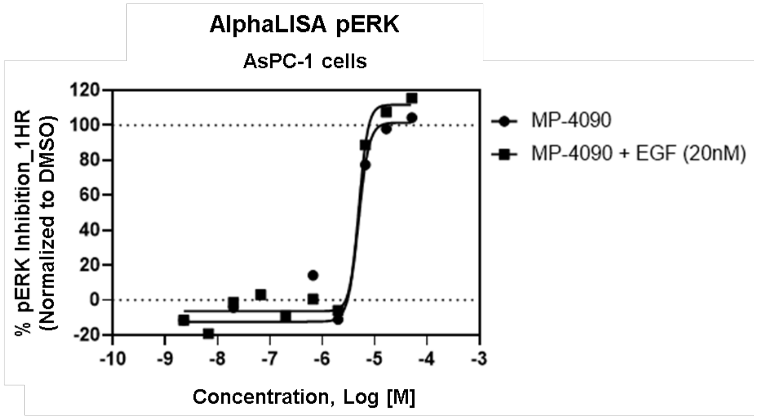


S9B)

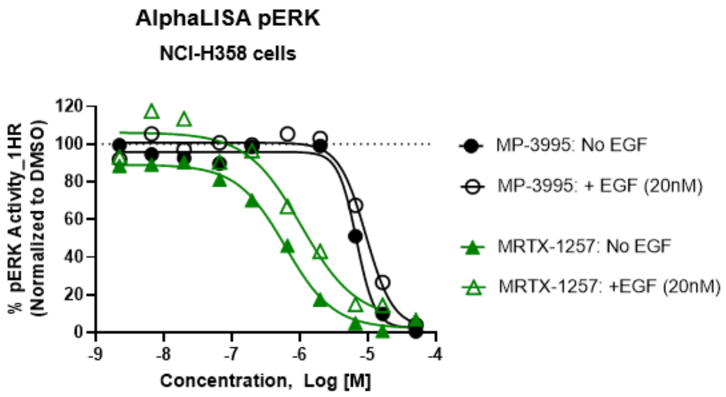


**Figure S9:** Cellular pERK activity of A) **MP-4090** on AsPC-1 (Homozygous KRAS$^{G12D}$) cells and B) **MP-3995** on NCI-H358 (Heterozygous KRAS$^{G12C}$) cells are not shifted by EGF treatment. In contrast, 2-fold potency shift was expected with **MRTX-1257** (KRAS$^{G12C}$ GDP state inhibitor) in the presence of 20nM EGF stimulation.

**Table S1**. Alanine scan on peptide Ac-K($N_3$)-RR-cyclo(c(methylene)PLYISYDPVC)-RR-$NH_2$ (**MP-1687**).

| Compound | Modification | $KRAS^{G12D}$ GDP TR-FRET $EC_{50}$ (nM) |
|---|---|---|
| **MP-1687** | none | 60 |
| **5** | Pro6Ala | >48080 |
| **6** | Leu7Ala | >48080 |
| **7** | Tyr8Ala | 691 |
| **8** | Ile9Ala | >48080 |
| **9** | Ser10Ala | 1072 |
| **10** | Tyr11Ala | 2430 |
| **11** | Asp12Ala | >48080 |
| **12** | Pro13Ala | 427 |
| **13** | Val14Ala | 161 |

**Table S2:** Effects on binding energy on in-silico alanine scanning of residues in **MP-1687** in complex with $KRAS^{G12D}$-GMPPCP

| **Mutation** | **ΔΔG (kcal/mol)** |
|---|---|
| Pro6Ala | 5 ±0.7 |
| Leu7Ala | 7 ±0.9 |
| Tyr8Ala | 3 ±0.9 |
| Ile9Ala | 6 ±0.7 |
| Ser10Ala | 3 ±0.5 |
| Try11Ala | 5 ±1.2 |
| Asp12Ala | 9 ±1.3 |
| Pro13Ala | 0.9 ±0.6 |
| Val14Ala | 0.5 ±0.2 |

**Table S3:** Cell culture conditions and seeding densities for 5 day CellTiter-Glo® cell viability assay on extended 16 cell panel.

| **Cell Line** | **Vendor** | **Catalog number** | **Description** | **Complete growth medium** | **Seeding density (cells/well)** |
|---|---|---|---|---|---|
| A375 | ATCC | CRL-1619 | skin, malignant melanoma | DMEM+10%FBS | 250 |
| A427 | ATCC | HTB-53 | lung, carcinoma | EMEM + 10%FBS | 400 |
| AsPC-1 | ATCC | CRL-1682 | pancreas, adenocarcinoma | RPMI1640+10%FBS | 900 |
| Capan-1 | ATCC | HTB-79 | pancreas, adenocarcinoma | IMDM+20%FBS | 700 |
| CFPAC-1 | ATCC | CRL-1918 | pancreas, ductal adenocarcinoma; cystic fibrosis | IMDM+10%FBS | 600 |
| HEK293 | ATCC | CRL-1573 | kidney, embryonic | EMEM+10%FBS | 600 |
| HPAF-II | ATCC | CRL-1997 | pancreas, adenocarcinoma | EMEM+10%FBS | 800 |
| MiaPaCa-2 | ATCC | CRL-1420 | pancreas, pancreatic Carcinoma | DMEM+10%FBS | 600 |
| NCI-H2122 | ATCC | CRL-5985 | lung, non-small cell lung cancer | RPMI1640+10%FBS | 400 |
| NCI-H358 | ATCC | CRL-5807 | lung, non-small cell lung cancer,bronchioalveolar carcinoma | RPMI1640+10%FBS | 500 |
| NCI-H727 | ATCC | CRL-5815 | lung, bronchus cancer | RPMI1640+10%FBS | 750 |
| Panc 08.13 | ATCC | CRL-2551 | pancreas, adenocarcinoma | RPMI1640 + 20 Units/ml human recombinant insulin +10%FBS | 900 |
| Panc 10.05 | ATCC | CRL-2547 | pancreas, adenocarcinoma | RPMI1640 + 20 Units/ml human recombinant insulin +10%FBS | 800 |
| SK-CO-1 | ATCC | HTB-39 | colon, colorectal adenocarcinoma | EMEM+10%FBS | 300 |
| SK-HEP-1 | ATCC | HTB-52 | liver, adenocarcinoma | EMEM+10%FBS | 600 |
| SW480 | ATCC | CCL-228 | colon, adenocarcinoma | Leibovitz's L-15 Medium+10%FBS | 350 |

**Table S4**. List of reported compounds and purity and characterization data.

| Compd. | Sequence | Method | Purity (%) | Formula | MW | Exact mass | Observed mass |
|---|---|---|---|---|---|---|---|
| **1** | Ac-K($N_3$)-RRRR-cyclo(DPLYISYDPV-Dap)-RRRR-$NH_2$ | D | 92.2 | C115H193N49O27 | 2694.13 | 2692.52 | 899.1 [M-3H]/3 |
| **2** | Ac-K($N_3$)-RRRR-cyclo(dPLYISYDPV-Dap)-RRRR-$NH_2$ | D | 93.0 | C115H193N49O27 | 2694.13 | 2692.52 | 899.1 [M-3H]/3 |
| **3** | Ac-K($N_3$)-RRRR-cyclo(C(methylene)PLYISYDPVC)-RRRR-$NH_2$ | E | 90.2 | C115H194N48O26S2 | 2729.26 | 2727.48 | 546.8 [M+5H]/5 |
| **4** | Ac-K($N_3$)-RR-cyclo(c(methylene)PLYISYDPV-hC)-RR-$NH_2$ | B | 92 | C92H148N32O22S2 | 2118.53 | 1398.64 | 1399.3 [M+2H]/2 |
| **MP-6483** | Ac-K($N_3$)-RRRR-cyclo(c(methylene)PLYISYDPVC)-RRRR- $NH_2$ | E | 95.3 | C115H194N48O26S2 | 2729.26 | 2727.48 | 569.5 [M-TFA-4H]/5 |
| **MP-1687** | Ac-K($N_3$)-RR-cyclo(c(methylene)PLYISYDPVC)-RR- $NH_2$ | E | 96.9 | C91H146N32O22S2 | 2104.51 | 2103.07 | 527.0 [M-4H]/4 |
| **S1** | Ac-RR-cyclo(c(methylene)ALYISYDPVC)-RR- $NH_2$ | F | 95.7 | C83H134N28O21S2 | 1924.3 | 1922.97 | 642.2 [M-3H]/3 |
| **S2** | Ac-RR-cyclo(c(methylene)PAYISYDPVC)-RR-$NH_2$ | F | 90.3 | C82H130N28O21S2 | 1908.25 | 1906.94 | 636.8 [M-3H]/3 |
| **S3** | Ac-RR-cyclo(c(methylene)PLAISYDPVC)-RR-$NH_2$ | F | 95.0 | C79H132N28O20S2 | 1858.24 | 1856.96 | 465.4 [M-4H]/4 |
| **S4** | Ac-RR-cyclo(c(methylene)PLYASYDPVC)-RR-$NH_2$ | F | 94.2 | C82H130N28O21S2 | 1908.25 | 1906.94 | 636.9 [M-3H]/3 |
| **S5** | Ac-RR-cyclo(c(methylene)PLYIAYDPVC)-RR-$NH_2$ | F | 96.0 | C85H136N28O20S2 | 1934.34 | 1932.99 | 645.5 [M-3H]/3 |
| **S6** | Ac-RR-cyclo(c(methylene)PLYISADPVC)-RR-$NH_2$ | F | 95.9 | C79H132N28O20S2 | 1858.24 | 1856.96 | 620.2 [M-3H]/3 |
| **S7** | Ac-RR-cyclo(c(methylene)PLYISYAPVC)-RR-$NH_2$ | F | 96.5 | C84H136N28O19S2 | 1906.32 | 1905.00 | 477.4 [M-4H]/4 |
| **S8** | Ac-RR-cyclo(c(methylene)PLYISYDAVC)-RR-$NH_2$ | F | 94.8 | C83H134N28O21S2 | 1924.3 | 1922.97 | 642.2 [M-3H]/3 |
| **S9** | Ac-RR-cyclo(c(methylene)PLYISYDPAC)-RR-$NH_2$ | F | 96.9 | C83H132N28O21S2 | 1922.28 | 1920.96 | 641.5 [M-3H]/3 |

| | | | | | | | |
|---|---|---|---|---|---|---|---|
| **MP-4090** | Ac-K($N_3$)-rrrr-cyclo(c(methylene)PLYISYDPVC)-rrrr-$NH_2$ | F | 95.1 | C115H194N48O26S2 | 2729.26 | 2727.48 | 683.1 [M-4H]/4 |
| **MP-3995** | Ac-K($N_3$)-rrrr-cyclo(c(methylene)-PLYI-aMeS-YDPVC)-rrrr-$NH_2$ | A | 99 | C116H196N48O26S2 | 2743.29 | 2741.49 | 915.7 [M+3H]/3 |
| **MP-4956** | Ac-rrrr-cyclo(c(methylene)cplyisydpvc)-rrrr-$NH_2$ | F | 95.3 | C109H185N44O25S2 | 2575.09 | 2574.40 | 644.6 [M-4H]/4 |
| **MP-9657** | Ac-K($N_3$)-rrrr-cyclo(c(methylene)PLYiSYDPVC)-rrrr-$NH_2$ | F | 96.0 | C115H194N48O26S2 | 2729.26 | 2727.48 | 910.5 [M-3H]/3 |
| **MP-9658** | Ac-K($N_3$)-rrrr-cyclo(c(methylene)PLYISYdPVC)-rrrr-$NH_2$ | F | 93.3 | C115H194N48O26S2 | 2729.26 | 2727.48 | 910.5 [M-3H]/3 |
| **5** | Ac-K($N_3$)-RR-cyclo(c(methylene)-aMeP-LYISYDPVC)-RR-$NH_2$ | A | 90 | C92H148N32O22S2 | 2118.53 | 2117.09 | 1059.9 [M+2H]/2 |
| **6** | Ac-K($N_3$)-RR-cyclo(c(methylene)P-aMeL-YISYDPVC)-RR-$NH_2$ | A | 90 | C92H148N32O22S2 | 2118.53 | 2117.09 | 1059.6 [M+2H]/2 |
| **7** | Ac-K($N_3$)-RR-cyclo(c(methylene)PL-aMeF-ISYDPVC)-RR-$NH_2$ | A | 92 | C92H148N32O21S2 | 2102.54 | 2101.09 | 1051.7 [M+2H]/2 |
| **8** | Ac-K($N_3$)-RR-cyclo(c(methylene)PLYI-aMeS-YDPVC)-RR-$NH_2$ | A | 90 | C92H148N32O22S2 | 2118.53 | 2117.09 | 1060.0 [M+2H]/2 |
| **9** | Ac-K($N_3$)-RR-cyclo(c(methylene)PLYIS-aMeY-DPVC)-RR-$NH_2$ | A | 94 | C92H151N32O22S2 | 2118.53 | 2117.09 | 1059.9 [M+2H]/2 |
| **10** | Ac-K($N_3$)-RR-cyclo(c(methylene)PLYISY-aMeD-PVC)-RR-$NH_2$ | C | 94 | C92H148N32O22S2 | 2118.53 | 2117.09 | 1059.7 [M+2H]/2 |
| **11** | Ac-K($N_3$)-RR-cyclo(c(methylene)PLYISYD-aMeP-VC)-RR-$NH_2$ | A | 90 | C92H148N32O22S2 | 2118.53 | 2117.09 | 1059.7 [M+2H]/2 |
| **12** | Ac-K($N_3$)-RR-cyclo(c(methylene)PLYISYDP-aMeV-C)-RR-$NH_2$ | A | 96 | C92H151N32O22S2 | 2118.53 | 2117.09 | 1060.1 [M+2H]/2 |
| **13** | Ac-K($N_3$)-RR-cyclo(c(methylene)PLYISYDPV-aMeC)-RR-$NH_2$ | A | 84 | C92H148N32O22S2 | 2118.53 | 2117.09 | 1059.8 [M+2H]/2 |
| **14** | Ac-K($N_3$)-RRR-cyclo(c(methylene)PLYISYDPVC)-RRR-$NH_2$ | E | 93.8 | C103H170N40O24S2 | 2416.88 | 2415.28 | 844.5 [M-TFA-2H]/3 |
| **MP-9903** | Ac-K($N_3$)-R-cyclo(c(methylene)PLYISYDPVC)-R-$NH_2$ | F | 96.6 | C79H122N24O20S2 | 1792.13 | 1790.87 | 896.7 [M-2H]/2 |
| **15** | Ac-K($N_3$)-cyclo(c(methylene)PLYISYDPVC)-R-$NH_2$ | F | 94.9 | C73H110N20O19S2 | 1635.94 | 1634.77 | 818.4 [M-2H]/2 |

| | | | | | | | |
|---|---|---|---|---|---|---|---|
| **16** | Ac-K($N_3$)-cyclo(c(methylene)PLYISYDPVC)-$NH_2$ | A | 99 | C67H98N16O18S2 | 1479.75 | 1478.67 | 740.5 [M+2H]/2 |
| **MP-6139** | 5FAM-K($N_3$)-RRRR-cyclo(c(methylene)PLYISYDPVC)-RRRR-$NH_2$ | F | 91.2 | C134H202N48O31S2 | 3045.54 | 3043.51 | 508.5 [M-6H]/6 |
| **MP-8996** | 5FAM-rrrr-cyclo(c(methylene)PLYISYDPVC)-rrrr-NH2 | F | 92.8 | C128H192N44O30S2 | 2891.36 | 2889.43 | 723.6 [M-4H]/4 |
| **MP-3958** | Ac-K($N_3$)-RRRR-cyclo(c(methylene)PLYISYdPVC)-RRRR-$NH_2$ | F | 96.9 | C115H194N48O26S2 | 2729.26 | 2727.48 | 546.7 [M-5H]/5 |

5FAM = 5-Carboxyfluorescein; Ac = acetyl; aMeC = α-methyl-L-cysteine; aMeD = α-methyl-L-aspartic acid; aMeF = α-methyl-L-phenylalanine; aMeL = α-methyl-L-leucine; aMeP = α-methyl-L-proline; aMeS = α-methyl-L-serine; aMeV = α-methyl-L-valine; aMeY = α-methyl-L-tyrosine; c = D-Cys; d = D-Asp; Dap = L-2,3-diaminopropionic acid; K(N3) = *N*-ε-azido-L-lysine; hC = L-homocysteine; i = D-Ile; l = D-Leu ; p = D-Pro; r = D-Arg; s = D-Ser; v = D-Val; y = D-Tyr.

### Materials and Methods

#### General procedure for the synthesis of compounds in the Table S4

**General Information:**

Peptides in Table S4 were synthesized using standard solid phase synthesis using Fmoc/tert-Bu chemistry as exemplified in Chan, W. C.; White, P. D. "Fmoc Solid-Phase Synthesis: a Practical Approach", Oxford University Press, Oxford, **2000**; Steward, J.; Young, J. "Solid Phase Peptide Synthesis", Pierce Chemical Company, Rockford, **1984**.; N. L. Benoiton, "Chemistry of Peptide Synthesis", CRC Press, New York, **2006**; and Lloyd-Williams, P.; Albericio, F. "Chemical Approaches to the Synthesis of Peptides and Proteins", CRC Press, New York, **1997**.

The α-amino group of each amino acid was protected by a 9*H*-fluoren-9-ylmethoxycarbonyl group (Fmoc) during the coupling of the carboxylic acid of the amino acid with the free amino terminus of the peptide attached to the resin. To avoid any side reactions during the coupling steps performed in DMF, the reactive side-chains of amino acids also carried acid-labile protecting groups that effectively masked the reactive groups until treatment of the resin with acid during the cleavage of the peptide from the solid support. After completion of each coupling step, the Fmoc group of the just-attached amino acid was removed with piperidine or 4-methylpiperidine and the resin was thoroughly washed to prepare for the coupling of the subsequent Fmoc-protected amino acid derivative.

The side-chain protecting groups were: *tert*-butyl (*t*Bu) for L-Asp, D-Asp, α-Me-L-Ser, L-Ser, and L-Tyr; trityl (Trt) for L-Asn, L-Cys and D-Cys; tert-butoxy-carbonyl (Boc) for L-Dap; and, 2,2,4,6,7-pentamethyldihydrobenzofuran-5-sulfonyl (Pbf) for L-Arg and D-Arg.

Fmoc-protected amino acids were typically obtained from vendors such as Sigma-Aldrich, Novabiochem, Chem-Impex, Combi-Block.

**Protocol A:**

**Step 1. Linear peptide synthesis.** The peptide was synthesized using Fmoc/*t*-Bu chemistry on Rink Amide resin LL MBHA (NovaBioChem, 0.33 mmol/g) with a CEM Liberty automated microwave peptide synthesizer. The peptide sequence was synthesized on a 0.1 mmol scale. Typical reaction conditions were as follows: Fmoc deprotections were performed using 20% (v/v) piperidine in DMF (2 min at 90 °C). Reaction coupling conditions: double couplings with Fmoc-protected amino acid/*N*,*N'*-diisopropylcarbodiimide(DIC)/Oxyma Pure (5, 10, and 5 equiv respectively; 90 °C microwave assisted heating, 2 min or 4 min for difficult couplings such as α-Me amino acids, repeated twice). After the completion of linear peptide synthesis, a final acetylation step was performed using acetic anhydride (10% in DMF; 75 °C for 10 min).

**Step 2. Cleavage and Deprotection.** The linear resin-bound peptides were deprotected and cleaved from the solid support by treatment with a cleavage solution (TFA/DTT/Thioanisole/Phenol/$H_2O$ 87.5/2.5/5/2.5/2.5 or TFA/TIS/water 95/2.2/2.5, v/v, 8 mL) at 42 °C for 30 min using a Razor® peptide cleavage system from CEM Corporation. After filtration of the resin, crude linear peptides were precipitated from the TFA cleavage solution using cold TBME (40 mL) and collected by centrifugation (4000 rpm). The crude peptide was washed with cold ethyl ether or TBME (35 mL) and air dried. The residue was semi-purified by Teledyne ISCO flash chromatography (15.5 g HP C18 Aq column) with mobile phase A: water with 0.1% TFA and mobile phase B: ACN with 0.1% TFA, gradient 15-35% over 12 column volumes. Fractions containing the desired product were collected and lyophilized to give the linear product.

**Step 3. Peptide Cyclization.** The above linear peptide was dissolved in acetonitrile/DI water (1:1, v/v, 15 mL). $(NH_4)_2CO_3$ (0.2 M solution in water) was added to adjust the pH to ~8. 3,6-Dioxa-1,8-octanedithiol (DODT, 2 equiv) or DL-Dithiothreitol (DTT, 2 equiv) and diiodomethane (20 equiv) were added. Additional acetonitrile (2 mL) was added to make the reaction mixture homogeneous. The resulting reaction mixture was stirred at room temperature overnight. After the reaction was complete, the reaction solution was quenched by addition of TFA (100 µL), concentrated and lyophilized. The residue was purified by RP-HPLC on Waters XSelect Peptide CSH C18 OBD Prep column (130Å, 5 µm, column size 150 × 30 mm) using a Waters MS-Directed AutoPurification HPLC/MS system. Mobile phase: (A) 0.1 % TFA in HPLC water and (B) 0.1 % TFA in HPLC acetonitrile; flow rate: 50 mL/min; UV wavelength λ = 214 nm; gradient: 5% B over 17 min. The desired fractions were then lyophilized to provide the cyclized peptide as a white powder.

**Protocol B:**

**Step 1. Linear peptide synthesis.** The peptide was synthesized using Fmoc/*t*-Bu chemistry on Rink Amide resin LL MBHA (NovaBioChem, 0.33 mmol/g) with a CEM Liberty automated microwave peptide synthesizer. The peptide sequence was synthesized on a 0.1 mmol scale. Typical reaction conditions were as follows: Fmoc deprotections were performed using 20% (v/v) piperidine in DMF (2 min at 90 °C). Reaction coupling conditions: double couplings with Fmoc-protected amino acid/DIC/Oxyma Pure (5, 10, and 5 equiv respectively; 90 °C microwave assisted heating, 2 min or 4 min for difficult couplings such as α-Me amino acids, repeated twice). After the completion of linear peptide synthesis, a final acetylation step was performed using acetic anhydride (10% in DMF; 75 °C for 10 min).

**Step 2. Cleavage and Deprotection.** The linear resin-bound peptides were deprotected and cleaved from the solid support by treatment with a cleavage solution (TFA/DTT/Thioanisole/Phenol/$H_2O$ 87.5/2.5/5/2.5/2.5 or TFA/TIS/water 95/2.2/2.5, v/v, 8 mL) at 42 °C for 30 min using a Razor® peptide cleavage system from CEM Corporation. After filtration of the resin, crude linear peptides were precipitated from the TFA cleavage solution using cold TBME (40 mL) and collected by centrifugation (4000 rpm). The crude peptide was washed with cold ethyl ether or TBME (35 mL) and air dried. The residue was semi-purified by Teledyne ISCO flash chromatography (15.5 g HP C18 Aq column) with mobile phase A: water with 0.1% TFA and mobile phase B: ACN with 0.1% TFA, gradient 15-35% over 12 column volumes. Fractions containing the desired product were collected and lyophilized to give the linear product.

**Step 3. Peptide Cyclization.** The above linear peptide was dissolved in acetonitrile/DI water (1:1, v/v, 15 mL). $(NH_4)_2CO_3$ (0.2 M solution in water) was added to adjust the pH to ~8. 3,6- DL-Dithiothreitol (DTT, 2 equiv) and 1,2-dibromoethane (10 equiv) were added. Additional acetonitrile (2 mL) was added to make the reaction mixture homogeneous. The resulting reaction mixture was stirred at room temperature overnight. After the reaction was complete, the reaction solution was quenched by addition of TFA (100 µL), concentrated and lyophilized. The residue was purified by RP-HPLC on Waters XSelect Peptide CSH C18 OBD Prep column (130Å, 5 µm, column size 150 × 30 mm) using a Waters MS-Directed AutoPurification HPLC/MS system. Mobile phase: (A) 0.1 % TFA in HPLC water and (B) 0.1 % TFA in HPLC acetonitrile; flow rate: 50 mL/min; UV wavelength λ = 214 nm; gradient: 5% B over 17 min. The desired fractions were then lyophilized to provide the cyclized peptide as a white powder.

**Protocol C:**

**Step 1. Linear peptide synthesis.** The peptide was synthesized using Fmoc/*t*-Bu chemistry on Rink Amide resin LL (NovaBioChem, 0.33 mmol/g) with a Symphony® X synthesizer (Gyros Protein Technologies). The peptide sequence was synthesized on a 0.1 mmol scale at RT. Typical reaction conditions were as follows: Fmoc deprotections were performed using 20% (v/v) piperidine in DMF (4 mL, 3 min, 3 repeats). Reaction coupling conditions: double couplings with Fmoc-protected amino acid/HATU/NMM (5, 5, and 10 equiv respectively; 30 min or 60 min for difficult couplings such as α-Me amino acids, repeated twice). After the completion of linear peptide synthesis, a final acetylation step was performed using acetic anhydride (1.0 M) and DIPEA (2.0 M) in NMP (*N*-Methyl-2-pyrrolidone) at room temperature for 10 min.

**Step 2. Cleavage and Deprotection.** The linear resin-bound peptides were deprotected and cleaved from the solid support by treatment with TFA/DTT/Thioanisole/Phenol/$H_2O$ (87.5/2.5/5/2.5/2.5, v/v; 8 mL) at 42 °C for 30 min using a Razor® peptide cleavage system from CEM Corporation. After filtration of the resin, crude linear peptides were precipitated from the TFA cleavage solution using cold ethyl ether (40 mL) and collected by centrifugation (4000 rpm). The crude peptide was washed with cold ethyl ether (35 mL) and air dried. The residue was semi-purified by Teledyne ISCO flash chromatography (15.5 g HP C18 Aq column) with mobile phase A: water with 0.1% TFA and mobile phase B: ACN with 0.1% TFA. Fractions containing the desired product were collected and lyophilized to give the linear product.

**Step 3. Peptide Cyclization.** The above linear peptide was dissolved in acetonitrile/DI water (1:1, v/v, 15 mL). $(NH_4)_2CO_3$ (0.2 M solution in water) was added to adjust the pH to ~8. DODT (2 equiv) and diiodomethane (20 equiv) were added. Additional acetonitrile (2 mL) was added to make the reaction mixture homogeneous. The resulting reaction mixture was stirred at room temperature overnight. After the reaction was complete, the reaction solution was quenched by addition of TFA (100 µL), frozen and lyophilized. The residue was purified by RP-HPLC on Waters XSelect Peptide CSH C18 OBD Prep column (130Å, 5 µm, column size 150 × 30 mm) using a Waters MS-Directed AutoPurification HPLC/MS system. Mobile phase: (A) 0.1 % TFA in HPLC water and (B) 0.1 % TFA in HPLC acetonitrile; flow rate: 50 mL/min; UV wavelength λ = 214 nm; gradient: 5% B over 17 min. The desired fractions were then lyophilized to provide the cyclized peptide as a white powder.

**Protocol D:**

**Step 1. Linear peptide synthesis.** The peptide sequence was synthesized manually using standard solid phase synthesis using Fmoc/*t*-Bu chemistry. HATU with NMM were used as coupling agents to create the amide bond between the free amino terminus of the resin-bound protected peptide and the carboxylic acid of the Fmoc-protected amino acid. Unloaded Rink amide MBHA resin (100-200 mesh, 1% cross-linked polystyrene) was used for synthesis. All the amino acids were dissolved at a 0.2 M concentration in anhydrous DMF. Reactions were typically performed at the 0.3 mmol scale.

Every synthesis cycle included: (1) Fmoc amino acid deprotection: 20% piperidine in DMF (20 mL) was added into the resin. The mixture was kept at room temperature for 30 min while a stream of nitrogen was bubbled through it. The mixture was filtered, and the peptidyl resin was washed with DMF (5 × 20 mL); (2) Coupling (potentially repeated twice for difficult couplings) with Fmoc-protected amino acid/HATU/NMM (3, 2.85 and 6 equiv, respectively; room temperature; 1 h). Cycles of Fmoc deprotection and Fmoc-protected amino acid coupling were repeated with the desired monomers until the full linear peptide was formed. Final acetylation (capping) step was performed using acetic anhydride capping solution [Capping Solution: Acetic anhydride (6 mL) and NMM (10 mL) in DMF (84 mL)].

**Step 2. Orthogonal Deprotection and Macrolactamization.** The Allyl and Alloc protecting groups on the orthogonally protected side-chains were removed by using $Pd(PPh_3)_4$ (0.2 equiv) and phenylsilane (20 equiv) in DCM. The mixture was shaken for 4 h and the solution was drained. The resin was washed with 5% (w/v) sodium diethyldithiocarbamate solution and then washed successively with DMF (5 × 4 mL), DCM (5 × 4 mL) and DMF (2 × 4 mL).

The deprotected carboxylic acid and amine side-chains were coupled by using PyAOP (3 equiv), HOBt (10 equiv) and DIPEA (5 equiv) in DMF. It was shaken for 16 h at room temperature, and then the resin was washed with DMF (5 × 4 mL) and DCM (5 × 4 mL).

**Step 3. Cleavage and Deprotection.** The linear resin-bound peptide was deprotected and cleaved from the solid support by treatment with TFA/EDT/Thioanisole/Phenol/$H_2O$ (87.5/2.5/5/2.5/2.5, v/v, 30 mL) at room temperature for 2.5 h. After filtration of the resin, crude linear peptides were precipitated from the TFA cleavage solution using cold diethyl ether (270 mL) and collected by centrifugation (4000 rpm). The precipitate was washed with cold diethyl ether (2 × 270 mL). The crude was dried under vacuum overnight to give the crude deprotected linear peptide as a solid.

The residue was purified by RP-HPLC on Phenomenex Luna C18 column (100Å, 10 µm, column size 200 × 25 mm). Mobile phase: (A) 0.1 % TFA in HPLC water and (B) 0.1 % TFA in 80%ACN+20%$H_2O$; flow rate: 50 mL/min; UV wavelength λ = 220 nm; gradient: 30% B over 60 min. The desired fractions were then lyophilized to provide the cyclized peptide as a white powder.

**Protocol E:**

**Step 1. Linear peptide synthesis.** The peptide sequence was synthesized manually using standard solid phase synthesis using Fmoc/*t*-Bu chemistry. HATU with NMM were used as coupling agents to create the amide bond between the free amino terminus of the resin-bound protected peptide and the carboxylic acid of the Fmoc-protected amino acid. Unloaded Rink amide MBHA resin (100-200 mesh, 1% cross-linked polystyrene) was used for synthesis. All the amino acids were dissolved at a 0.2 M concentration in anhydrous DMF. Reactions were typically performed at the 0.3 mmol scale.

Every synthesis cycle included: (1) Fmoc amino acid deprotection: 20% piperidine in DMF (20 mL) was added into the resin. The mixture was kept at room temperature for 30 min while a stream of nitrogen was bubbled through it. The mixture was filtered, and the peptidyl resin was washed with DMF (5 × 20 mL); (2) Coupling (potentially repeated twice for difficult couplings) with Fmoc-protected amino acid/HATU/NMM (3, 2.85 and 6 equiv, respectively; room temperature; 1 h). Cycles of Fmoc deprotection and Fmoc-protected amino acid coupling were repeated with the desired monomers until the full linear peptide was formed. Final acetylation (capping) step was performed using acetic anhydride capping solution [Capping Solution: Acetic anhydride (6 mL) and NMM (10 mL) in DMF (84 mL)].

**Step 2. Cleavage and Deprotection.** The linear resin-bound peptide (~1.7 g of dry resin) was deprotected and cleaved from the solid support by treatment with TFA/EDT/Thioanisole/Phenol/$H_2O$ (87.5/2.5/5/2.5/2.5, v/v, 30 mL) at room temperature for 2.5 h. After filtration of the resin, crude linear peptides were precipitated from the TFA cleavage solution using cold diethyl ether (270 mL) and collected by centrifugation (4000 rpm). The precipitate was washed with cold diethyl ether (2 × 270 mL). The crude was dried under vacuum overnight to give the crude deprotected linear peptide as a solid (~450 mg).

**Step 3. Intermediate Purification.** The crude linear peptide was purified by RP-HPLC on Phenomenex Luna C18 column (100Å, 10 µm, column size 200 × 25 mm). Mobile phase: (A) 0.1 % TFA in HPLC water and (B) 0.1 % TFA in 80%ACN+20%$H_2O$; flow rate: 15 mL/min; UV wavelength λ = 220 nm; gradient: 30% B over 60 min. The desired fractions were then lyophilized to provide the linear peptide as a white powder.

**Step 4. Peptide Cyclization.** A portion of the above linear peptide was dissolved in phosphate buffered saline (40 mL, pH=7.2) and acetonitrile (40 mL). DTT (2 equiv) and diiodomethane (20 equiv) were added. The resulting reaction mixture was stirred at room temperature overnight. After the reaction was complete, the reaction solution was quenched by addition of TFA (100 µL) and lyophilized. The residue was purified by RP-HPLC on Phenomenex Luna C18 column (100Å, 10 µm, column size 200 × 25 mm). Mobile phase: (A) 0.1 % TFA in HPLC water and (B) 0.1 % TFA in 80%ACN+20%$H_2O$; flow rate: 50 mL/min; UV wavelength λ = 220 nm; gradient: 30% B over 60 min. The desired fractions were then lyophilized to provide the cyclized peptide as a white powder.

**Protocol F:**

**Step 1. Linear peptide synthesis.** The peptide sequence was synthesized manually using standard solid phase synthesis using Fmoc/*t*-Bu chemistry. HATU with NMM were used as coupling agents to create the amide bond between the free amino terminus of the resin-bound protected peptide and the carboxylic acid of the Fmoc-protected amino acid. Unloaded Rink amide MBHA resin (100-200 mesh, 1% cross-linked polystyrene) was used for synthesis. All the amino acids were dissolved at a 0.2 M concentration in anhydrous DMF. Reactions were typically performed at the 0.3 mmol scale.

Every synthesis cycle included: (1) Fmoc amino acid deprotection: 20% piperidine in DMF (20 mL) was added into the resin. The mixture was kept at room temperature for 30 min while a stream of nitrogen was bubbled through it. The mixture was filtered, and the peptidyl resin was washed with DMF (5 × 20 mL); (2) Coupling (potentially repeated twice for difficult couplings) with Fmoc-protected amino acid/HATU/NMM (3, 2.85 and

6 equiv, respectively; room temperature; 1 h). Cycles of Fmoc deprotection and Fmoc-protected amino acid coupling were repeated with the desired monomers until the full linear peptide was formed. Final acetylation (capping) step was performed using acetic anhydride capping solution [Capping Solution: Acetic anhydride (6 mL) and NMM (10 mL) in DMF (84 mL)].

**Step 2. Cleavage and Deprotection.** The linear resin-bound peptide (~1.7 g of dry resin) was deprotected and cleaved from the solid support by treatment with TFA/EDT/Thioanisole/Phenol/$H_2O$ (87.5/2.5/5/2.5/2.5, v/v, 30 mL) at room temperature for 2.5 h. After filtration of the resin, crude linear peptides were precipitated from the TFA cleavage solution using cold diethyl ether (270 mL) and collected by centrifugation (4000 rpm). The precipitate was washed with cold diethyl ether (2 × 270 mL). The crude was dried under vacuum overnight to give the crude deprotected linear peptide as a solid (~450 mg).

**Step 3. Intermediate Purification.** The crude linear peptide was purified by RP-HPLC on Phenomenex Luna C18 column (100Å, 10 µm, column size 200 × 25 mm). Mobile phase: (A) 0.1 % TFA in HPLC water and (B) 0.1 % TFA in 80%ACN+20%$H_2O$; flow rate: 15 mL/min; UV wavelength λ = 220 nm; gradient: 30% B over 60 min. The desired fractions were then lyophilized to provide the linear peptide as a white powder.

**Step 4. Peptide Cyclization.** A portion of the above linear peptide was dissolved in acetonitrile/DI water (1:1, v/v, 15 mL). DODT (2 equiv) was added and the mixture was stirred at room temperature for 30 min. $(NH_4)_2CO_3$ (0.2 M solution in water) was added to adjust the pH to ~8. DODT (2 equiv) and diiodomethane (20 equiv) were added. The reaction mixture was stirred at room temperature overnight. After the reaction was complete, the reaction solution was quenched by addition of TFA (100 µL), frozen and lyophilized. The residue was purified by RP-HPLC on Phenomenex Luna C18 column (100Å, 10 µm, column size 200 × 25 mm). Mobile phase: (A) 0.1 % TFA in HPLC water and (B) 0.1 % TFA in 80%ACN+20%$H_2O$; flow rate: 50 mL/min; UV wavelength λ = 220 nm; gradient: 30% B over 60 min. The desired fractions were then lyophilized to provide the cyclized peptide as a white powder.

**TR-FRET tracer**

KRAS$^{G12D}$ (GDP) binding was measured by a TR-FRET competitive binding assay using a 5-FAM-labeled analog of Pro13Lys **MP-1687** with the following structure:

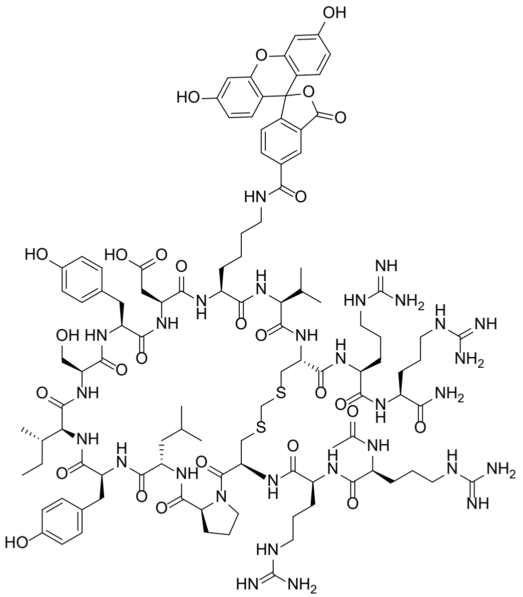


**Step 1. Solid Phase Synthesis.** Peptide was synthesized manually using standard solid phase synthesis using Fmoc/*t*-Bu chemistry as previously reported. HATU with NMM were used as coupling agents to create the amide bond between the free amino terminus of the resin-bound protected peptide and the carboxylic acid of the Fmoc-protected amino acid. The side chain protecting groups were: *tert*-butyl (*t*Bu) for L-Asp, L-Ser and L-Tyr; trityl (Trt) for L-Cys and D-Cys; 1-(4,4-dimethyl-2,6-dioxocyclohex-1-ylidene)ethyl (Dde) for L-Lys; 2,2,4,6,7-pentamethyldihydrobenzofuran-5-sulfonyl (Pbf) for L-Arg. Knorr Amide MBHA Resin (0.27 mmol/g loading) was used for synthesis. All the amino acids were dissolved at a 0.2 M concentration in anhydrous DMF. The reaction was performed at the 0.2 mmol scale.

Every synthesis cycle included: (1) Fmoc amino acid deprotection: 20% Piperidine in DMF (20 mL) was added into the resin. The mixture was kept at room temperature for 30 min while a stream of nitrogen was bubbled through it. The mixture was filtered, and the peptidyl resin was washed with DMF (5 × 20 mL); (2) Coupling (potentially repeated twice for difficult couplings) with Fmoc protected amino acid/HATU/NMM (3, 2.85 and 6 equiv, respectively; room temperature; 1 h). Cycles of Fmoc deprotection and Fmoc-protected amino acid coupling were repeated with the desired monomers until the full linear peptide was formed. Final acylation (capping) step was performed using acetic anhydride capping solution [Capping Solution: Acetic anhydride (6 mL) and NMM (10 mL) in DMF (84 mL)].

**Step 2. Orthogonal Deprotection.** Deprotection of the Lys side chain Dde: 4% hydrazine hydrate in DMF (15 mL) was added to the peptidyl resin. The mixture was kept at room temperature for 30 min while a stream of nitrogen was bubbled through it. The mixture was filtered, and the peptidyl resin was washed with DMF (5 × 20 mL). Then peptidyl resin was washed with MeOH (2 × 20 mL), DCM (2 × 20 mL) and MeOH (2 × 20 mL). The resin was dried under vacuum.

**Step 3. Coupling of 5-Carboxyfluorescein (5-FAM).** After selective deprotection of the Lys side chain, the peptidyl resin was treated with a mixture of 5-FAM (0.6 mmol, 3 equiv), HATU (0.57 mmol, 2.85 equiv) and NMM (1.2 mmol, 6 equiv) in DMF (2 mL). The suspension was shaken at room temperature for 1 h while a stream of nitrogen was bubbled through it. The peptidyl resin was washed with DMF (5 × 20 mL), MeOH (2 × 20 mL), DCM (2 × 20 mL) and MeOH (2 × 20 mL). The resin was dried under vacuum.

**Step 4. Cleavage and Deprotection.** The resin-bound peptide was deprotected and cleaved from the solid support by treatment with TFA/EDT/Thioanisole/Phenol/$H_2O$ (87.5/2.5/5/2.5/2.5, v/v, 34 mL) at room temperature for 2.5 h. After filtration of the resin, crude linear peptides were precipitated from the TFA cleavage solution using cold diethyl ether (150 mL) and collected by centrifugation (4000 rpm). The precipitate was washed with cold diethyl ether (2 × 170 mL). The crude was dried under vacuum overnight to give the crude deprotected cyclic peptide as a solid (475 mg).

**Step 5. Linear Peptide Purification.** Purification of the crude linear 5FAM-labeled peptide was performed by preparative reversed-phase high performance liquid chromatography (RP-HPLC) on Phenomenex Luna C18(2) column (100Å, 10 µm, column size 200 × 21.2 mm using Waters 4000 system. Mobile phase: (A) 0.1% TFA in water and (B) 0.1%TFA in 80%ACN+20%$H_2O$; flow rate: 15 mL/min; UV wavelength λ = 220 nm; gradient: 27–57% B over 60 min. UV absorbing fractions containing the target m/z ions were collected and the fractions containing product were confirmed by LC/MS.

**Step 6. Methylene Bridge Insertion.** The linear peptide (130 mg, 0.056 mmol) was dissolved in a 1:1 mixture of water/ACN (30 mL). DODT (0.112 mmol, 2 equiv) was added and the mixture was stirred at 30°C for 30 min. pH was adjusted to ~8 by addition of 5.8 mL of aqueous 0.2 M ammonium carbonate solution. Diiodomethane (1.12 mmol, 20 equiv) and DODT (0.112 mmol, 2 equiv) were added. The mixture was shaken at 30°C overnight.

The final crude 5-FAM-labeled peptide was purified by preparative reversed-phase high performance liquid chromatography (RP-HPLC) on Phenomenex Luna C18(2) column (100Å, 10 µm, column size 200 × 21.2 mm using Waters 4000 system. Mobile phase: (A) 0.1% TFA in water and (B) 0.1%TFA in 80% ACN+20% $H_2O$; flow rate: 15 mL/min; UV wavelength λ = 220 nm; gradient: 22–52% B over 60 min. UV absorbing fractions containing the target m/z ions were collected and the fractions containing product were confirmed by LC/MS. The solution was collected and lyophilized to give the final tracer compound as a solid 11.0 mg with the purity of 95.3%.

Purity of fractions was confirmed by UPLC, which was measured by a reverse phase Hewlett Packard 1100 UPLC-MS system. Column: Sepax GP-C18 Column (120Å, 5 µm, column size 150 × 4.6 mm). Mobile phase: (A) 0.1 % TFA in water and (B) 0.09 % TFA in 80%ACN+20%$H_2O$; gradient: 32–42% B in 20 min; flow rate: 1 mL/min; UV wavelength λ = 220 nm. The peptide was characterized by electrospray mass spectrometry on an Agilent 1260-6120 Quadrupole LC/MS (MW expected: 2339.7 Da; MW found: 585.8 ($[M+4H]^{4+}$).

**Protein Production:** The KRAS amino acid sequence used was derived from UniProt reference P01116-2 (KRAS4B isoform) RASK_HUMAN Isoform 2B of GTPase KRAS, Homo sapiens having a G12D mutation (VAR_016026). It comprised the following sequence (AAs 1 to 169 plus a 4xGly motif appended to the N-terminus for sortase tagging):

GGGGMTEYKLVVVGA**D**GVGKSALTIQLIQNHFVDEYDPTIEDSYRKQVVIDGETCLLDILDTAGQEEYSAMRDQYMRTGEGFLCVFAINNTKSFEDIHHYREQIKRVKDS
EDVPMVLVGNKCDLPSRTVDTKQAQDLARSYGIPFIETSAKTRQGVDDAFYTLVREIRKHKEK

Expression and purification of GDP and GMPPNP-loaded KRAS$^{G12D}$ were carried out. His -SUMO-tagged human KRAS resides 1-169 was expressed in *E. coli* Rosetta 2 (DE3). Protein expression was induced by 1 mM IPTG overnight at 18 °C. Cells were lysed in 40 mM Tris pH 7.8, 150 mM NaCl, 0.5% Triton x-100, 5 mM BME, 10 mM imidazole, protease inhibitors EDTA-free tablets (1/50mL). Clarified lysate was loaded on Ni-NTA FF prepacked columns (Qiagen) and washed with 10 column volumes (CV) of wash buffer (40 mM Tris pH 7.8, 150 mM NaCl, 5 mM BME, 10 mM imidazole). Protein was eluted by buffer containing 40 mM Tris pH 7.8, 150 mM NaCl, 5 mM BME, 300 mM imidazole. The resulting protein was tag removed by SUMO protease in dialysis buffer (40 mM Tris pH 7.8, 150 mM NaCl, 1 mM DTT, 10 mM Imidazole) overnight at 4 °C, followed by subtractive Ni-NTA purification to remove the tagged protein. KRAS protein was *N*-terminally biotinylated using SrtA and biotin-Sortag peptide [biotin-KGGGLPETGG-OHse(Ac)-amide]. The reaction was carried out at room temperature for 5 h, followed by Ni subtraction to remove His-tagged SrtA and SoftLink™ Avidin Resin purification to isolate the biotin-labeled K-Ras.

**GDP Nucleotide Loading:** KRAS$^{G12D}$ protein was treated with 5 mM EDTA and 5 mM GDP for a minimum of 2 h on ice. The reaction was stopped by addition of 40 mM $MgCl_2$ and incubate on ice for 1 h. The reaction was stopped by addition of 40 mM $MgCl_2$ and incubate on ice for 1 h. The resulting proteins were further purified and buffer-exchanged on a Superdex 75 26/60 size exclusion column preequilibrated in 25 mM HEPES pH 7.5, 150 mM NaCl, 2 $MgCl_2$, 2 mM TCEP. The nucleotide loading efficiency was analyzed by HPLC. Specifically, 5 µL of the protein or nucleotide at 100 µM were injected on a RP-C18 analytical column. Nucleotides were separated through an isocratic elution in 100 mM potassium phosphate, pH 6.5, 10 mM tetrabutylammonium bromide, 0.2 mM $NaN_3$, and 3% acetonitrile. The runs are monitored at 254 nm and compared to GDP standard.

**X-ray crystallography:** KRAS(1-168, G12D) loaded with GMPPCP was provided by Evotec at a concentration of 10mg/ml. The complex with **MP-9903** was prepared by incubating 3-fold molar excesses of the peptide on ice for 2h prior to the crystallization experiments. The complex was crystallized from a reservoir solution containing 0.1 M Bis-Tris (pH 6.5) and 25% (v/v) PEG 4000 at 18°C via the sitting-drop vapor diffusion method. Crystals were harvested and cryoprotected with an additional 10% glycerol. X-ray diffraction data were collected at IMCA beamline ID17 and processed using the Globalphasing [1] and CCP4 [2] packages. Structures were solved by molecular replacement using 5XCO [3] and refined using BUSTER [1] and Coot [4]. Peptides were modeled using standard amino acid residue definitions or restrains generated using Flynn [5]. Coordinates and experimental data are available from the Protein Data Bank with the accession code **7ROV**.

**Surface Plasmon Resonance:** SPR experiments were performed with Biacore T100 (GE Healthcare) at 25 °C. The site-specific mono-biotinylated proteins were prepared by sortase-mediated ligation. SPR buffer consisted of 50 mM Tris pH 7.4, 150 mM NaCl, 2 mM $MgCl_2$, 10 µM GDP, 0.05% Tween 20, and 3% DMSO. For reducing condition, 1 mM DTT was included in the SPR buffer and peptide solution. The CM5 chip was first conditioned with 100 mM HCl, followed by 0.5% SDS, 50 mM NaOH and then water, all performed twice with 6 sec injection at a flow rate of 100 µL/min. With the flow rate set to 10 µL/min, streptavidin (S4762, Sigma-Aldrich) was immobilized on the conditioned chip through amine coupling as described in the Biacore manual. Excess protein was removed by 30 sec injection of the wash solution (50 mM NaOH + 1 M NaCl) for at least 8 times. The immobilized levels are typically 4000 RU. The biotinylated $KRAS^{G12D}$ (GDP) protein was captured to the immobilized streptavidin to a level of ~300 RU. Flow cell consisted of only streptavidin was used as the reference surface. **KRpep-2d** (five concentrations: 0.6 nM, 1.9 nM, 5.6 nM, 16.7 nM, 50 nM) dissolved in the SPR buffer were injected sequentially in single-cycle kinetic format without regeneration (contact time 180 sec, dissociation time 300 sec, flow rate 30 mL/min). The dissociation was monitored for 300 sec. After the run, responses from the target protein surface are transformed by (i) correcting with DMSO calibration curve, (ii) subtracting the responses obtained from the reference surface, and (iii) subtracting the responses of buffer injections from those of peptide injections. The last step is known as double referencing which corrects the systematic artefacts. The resulting responses were subjected to kinetic analysis by fitting with 1:1 binding model.

**Time-Resolved Fluorescence/Förster Resonance Energy Transfer (TR-FRET) Assay using KRAS$^{G12D}$ Loaded with GDP:** Macrocyclic peptide binding affinities to KRAS$^{G12D}$ (GDP) were characterized using a TR-FRET–based competitive binding assay. A master mixture of 30 nM KRAS$^{G12D}$ (GDP), 5 nM Terbium-Streptavidin and 500 nM fluorescently labeled tracer (MP-6139) was prepared. 10 µL of the master mixture was added to each well of the 384-well, flat bottom, non-binding, black microplate (Greiner 784900) for the respective assay. Compound dose-response titrations were prepared, and appropriate amounts of compounds were dispensed, using Echo 550 liquid handler, into the respective 384-well black microplate, containing the master mixture. The assay plate is sealed with an aluminum foil and placed on a shaker at ambient temperature for 5 min. The plate was then centrifuged at 1000g for 3 min, then incubated in the dark for 18 h at ambient temperature. Assay plates were read at ambient temperature on the Tecan M1000, with excitation at 340 nm and emission at 495 and 520 nm. Dose response curves and $EC_{50}$ were analyzed using a 4-parameter logistic equation in GraphPad Prism software (GraphPad, San Diego, CA).

**KRAS/RBD PPI assay:** TR-FRET based RBD assay was carried out with a final concentration of 2.5 nM Tb-Streptavidin labeled KRAS$^{G12D}$ (GMPPNP), 10 nM GST-RBD and 1 ug/ml anti-GST-d2 (Cisbo) in RBD buffer (20 mM HEPES, pH 7.4, 150 mM NaCl, 10 mM MgCl2, 0.01%Tween). The assay volume was 10 µl in a 384-well plate (Greiner 784900). Inhibitors/DMSO vehicle were dispensed using Echo 550 liquid handler, into plate containing the reaction mixture. Plates were sealed and incubated at room temperature for 1h. TR-FRET signal was detected using EnVision plate reader with an excitation filter at 340 nm and emission filters for Tb at 615 and d2 at 665 nm, respectively. Dose response curves and $EC_{50}$ were analyzed using a 4-parameter logistic equation in GraphPad Prism software (GraphPad, San Diego, CA).

**SOS-mediated nucleotide exchange assay:** SOS-mediated nucleotide exchange assay was performed as previously reported [6] with slight modifications. Biotin-labeled $KRAS^{G12D}$ protein was treated with EDTA buffer (20 mM HEPES, pH 7.4, 50 mM NaCl, 10 mM EDTA, and 0.01% (w/v) Tween20) for 1 hour at room temperature. The resulting proteins were incubated with 10X excess of BODIPY-GDP in SOS assay buffer (20 mM HEPES, pH 7.4, 50 mM NaCl, 10 mM MgCl2, 0.01%(w/v) Tween20) for 6 hours at room temperature. A master mixture containing 3 nM K-Ras, 30 nM BODIPY-GDP and 0.5 nM Tb-Streptavidin was prepared. Inhibitors/DMSO vehicle were dispensed to a dry plate (Corning3820) using Echo 550 liquid handler; subsequently, 3 µl of master mixture and 3 µl of SOS assay buffer was added to the plate and sealed. Sealed plates were incubated at room temperature for 60 min prior to the addition of 3 µl 120 nM SOS1(618-1099 a.a.) and 9 mM GTP premix. Plates were read after 60 min incubation at room temperature using EnVision plate reader with excitation at 340 nm and emission at 495 and 520 nm. Dose response curves and $EC_{50}$ were analyzed using a 4-parameter logistic equation in GraphPad Prism software (GraphPad, San Diego, CA).

**Procedure for Cellular Phospho-ERK Assay in AsPC-1 Cells (Alpha Screen):** Cellular KRAS inhibitory activity was evaluated by phosphorylation levels of ERK1/2 in AsPC-1 (homozygous $KRAS^{G12D}$) cells. AsPC-1 cells (ATCC® CRL-1682™) were cultured in T175 flask in growth medium (RPMI 1640 Medium, GlutaMAX™ Supplement, HEPES (Gibco 72400-047) supplemented with 10% fetal bovine serum (Hyclone SH30071.03) and 1X Penicillin/Streptomycin (Gibco 15140-122). The cells were harvested in seeding medium (RPMI 1640 Medium, no phenol red (Gibco 11835-030) supplemented with 10% fetal bovine serum (Hyclone SH30071.03), 25 mM HEPES (Gibco 15630-080) and 1X Penicillin/Streptomycin (Gibco 15140-122) after 5 min of 0.25% Trypsin-EDTA (Gibco 25200-056) digestion and were seeded in 384-well tissue culture treated plate (Greiner 781091) at a density of 15,000 cells/20µL/well, and incubated at 37°C, 5% $CO_2$ overnight. Prior to dosing, seeding medium was removed using the BlueCatBio Bluewasher system and replaced with 20 µL of assay medium without fetal bovine serum (RPMI 1640 Medium, no phenol red (Gibco 11835-030) supplemented with 25 mM HEPES (Gibco 15630-080) and 1X Penicillin/Streptomycin (Gibco 15140-122). For assays performed in the presence of 10% serum, seeding medium was used instead of assay medium. The compound dose-response titrations were prepared, and appropriate amounts of compounds were dispensed into the 384-well cell culture assay plate using the Echo 550 liquid handler. 25 µL assay or seeding medium was added to achieve a final assay volume of 45 µL. Assay plate was incubated at 37°C, 5% $CO_2$ for the indicated length of time. After treatment, 25 µL medium was removed and transferred to an empty 384-well tissue culture treated plate (Greiner 781091) for the CytoTox-ONE™ Homogeneous Membrane Integrity (LDH) Assay. Remaining medium was removed from the plate using the BlueCatBio Bluewasher system, and cells were washed once with 25µL 1 × DPBS (Gibco 14190-144). Cells were lysed in 20 µL 1 × lysis buffer from Alpha *SureFire® Ultra*™ Multiplex pERK and total ERK assay kit (PerkinElmer MPSU-PTERK) containing EDTA-free Protease inhibitor cocktail (Roche 11836170001) at ambient temperature with constant shaking at 300 rpm for 10-15 min. The cell lysates were mixed for 10 cycles using the Agilent Bravo 384ST liquid handler system before 10 µL was transferred to OptiPlate-384 plate (PerkinElmer 6007680). Phosphorylated ERK and total ERK levels were detected by Alpha SureFire Ultra Multiplex pERK kit (PerkinElmer MPSU-PTERK) using 5 µL acceptor bead mix and 5 µL donor bead mix, both prepared following the manufacturer's protocol. Plates were sealed using aluminum sealing tape (Costar 07-200-683) during incubation at ambient temperature with constant shaking at 300 rpm for 1 h (both acceptor and donor). Assay plates were read on a Envision Xcite Multilabel Reader (PerkinElmer 1040900) at ambient temperature, with emission at 535 nm (Total ERK) and emission at 615 nm (Phospho ERK). Ratio of pERK vs total ERK in each well was used as the final readout. Dose response curves and $EC_{50}$ were analyzed using a 4-parameter logistic equation in GraphPad Prism software (GraphPad, San Diego, CA).

**Procedure for Cellular Phospho-ERK Assay in other cell lines:** Procedure for the Alpha screen assay on other cell lines were performed according to AsPC-1 protocol, with the following change in media and cell seeding densities: NCI-H2122 (ATCC® CRL-5985$^{TM}$), NCI-H358 (ATCC® CRL-5807$^{TM}$), and Panc 08.13 (ATCC® CRL-2551$^{TM}$) cells were cultured in T175 flask in growth medium (RPMI 1640 Medium, GlutaMAX™ Supplement, HEPES (Gibco 72400-047) supplemented with 10% fetal bovine serum (Hyclone SH30071.03) and 1X Penicillin/Streptomycin (Gibco 15140-122). Panc 08.13 was further supplemented with 20µg/mL of insulin, human recombinant (Gibco 12585-014). SK-CO-1 (ATCC® HTB-39$^{TM}$) and HEK293 (ATCC® CRL-1573$^{TM}$) cells were cultured in (MEM Medium, GlutaMAX™ Supplement (Gibco 42360-032) supplemented with 10% fetal bovine serum (Hyclone SH30071.03) and 1X Penicillin/Streptomycin (Gibco 15140-122). A-375 (ATCC® CRL-1619$^{TM}$) cells were cultured in (DMEM Medium, GlutaMAX™ Supplement (Gibco 10566-016) supplemented with 10% fetal bovine serum (Hyclone SH30071.03) and 1X Penicillin/Streptomycin (Gibco 15140-122). NCI-H2122, NCI-H358, SK-CO-1 and HEK293 cells were seeded at a density of 15,000 cells/20µL/well.

A375 cells were seeded at a density of 10,000 cells/20µL/well. For assays performed in the presence of EGF, cells were serum starved overnight prior to dosing, followed by 20nM of recombinant human EGF (Peprotech AF-100-15) stimulation for 10-15 minutes before cell lysates were harvested.

**CytoTox-ONE™ Homogeneous Membrane Integrity (LDH) Assay:** Transient damage to cellular membranes were measured by LDH release assay. At 1 and 18 h post dose, 25 µL medium was removed from the primary cell culture assay plate and transferred to an empty 384-well tissue culture treated plate (Greiner 781091) using the Agilent Bravo 384ST liquid handler system. CytoTox-ONE reaction mix was prepared from the CytoTox-ONE™ Homogeneous Membrane Integrity Assay Kit (Promega G7891) according to manufacturer's protocol. 25 µL of CytoTox-ONE reaction mix was added to the assay plate containing 25 µL medium, and plate was sealed using aluminum sealing tape (Costar 07-200-683). Assay plate was incubated at ambient temperature with constant shaking at 300 rpm for 45 min. Assay plates were read at ambient temperature on the Tecan M1000, with excitation at 560 nm and emission at 590 nm. Dose response curves and $EC_{50}$ were analyzed using a 4-parameter logistic equation in GraphPad Prism software (GraphPad, San Diego, CA).

**CellTiter-Glo® 2.0 Cell Viability (CTG) Assay:** Cell proliferation inhibition of compounds were determined using the CellTiter-Glo® 2.0 assay kit (Promega G9243). AsPC-1, NCI-H2122, NCI-H358, SK-CO-1 and A-375 cells were cultured in their respective growth medium containing 10% fetal bovine serum as according to the protocol described in the Alphascreen assay. AsPC-1 cells were seeded at a density of 1,000 cells/20µL/well into 384-well tissue culture treated plate (Greiner 781080). NCI-H2122, NCI-H358 and SK-CO-1 cells were seeded at a density of 500 cells/20µL/well. A-375 cells were seeded at a density of 300 cells/20µL/well. Assay plates were incubated at 37°C, 5% $CO_2$ overnight to allow cell attachment. Following day, compounds were dispensed into the assay plate using the Echo 550 liquid handler. 25 µL growth medium was added to achieve a final assay volume of 45 µL and assay plates were incubated at 37°C, 5% $CO_2$ for 120 h. At 120 h post dose, CellTiter-Glo assay was performed according to the manufacturer's protocol. 45 µL of CellTiter-Glo® 2.0 reagent was added to the assay plate containing 45 µL medium and incubated at room temperature for 30 minutes. Assay plates were read on a Envision Xcite Multilabel Reader (PerkinElmer 1040900) with luminescence at an integration time of 0.15 second per well. Dose response curves and $EC_{50}$ were analyzed using a 4-parameter logistic equation in GraphPad Prism software (GraphPad, San Diego, CA). AMG510 was prepared internally from published procedures.

**CellTiter-Glo® Cell Viability Assay (Extended cell panel):** Cell proliferation inhibition of compounds were performed on extended cell panels at fee-for-service at Shanghai ChemPartner Co., Ltd. Cell viabilities were determined using the CellTiter-Glo® assay kit (Promega G7558). Cells were cultured and seeded onto 384-well assay plate (Corning 3765) using the Thermo Scientific Multidrop, according to conditions described in Table S3. Assay plates were incubated at 37°C, 5% $CO_2$ overnight to allow cell attachment. Cells cultured using Leibovitz's L-15 medium were incubated in incubator without $CO_2$. Following day, compounds were dispensed into the assay plate using the Tecan HPD300 system and incubated at 37°C, 5% $CO_2$ for 120 h. At 120 h post dose, CellTiter-Glo assay was performed according to the manufacturer’s protocol. Assay plates were read on a Envision Reader with luminescence at an intergration time of 0.1 second per well. Staurosporine (Selleckchem S1421) was used as a positive control for the assay.

**Western Blotting:** Cells were seeded in 24-well poly-D-lysine coated plates and treated with the indicated peptides the next day. Following incubation, 100 µl of 1X Bolt™ LDS sample buffer supplemented with NuPAGE® sample reducing agent was added. Wells were scrapped using wide orifice tips and lysate was transferred into PCR-strip tubes and sonicated for 10 X 10 seconds in a chilled water bath sonicator (QSonica). 15 µl of protein extract was separated on 4-12% Bis-Tris plus gels, transferred onto nitrocellulose membranes using the Trans-Blot® Turbo™ semi-dry system (Bio-rad), and blocked for 1 hour at room temperature with tris-buffered saline (TBS) Odyssey blocking buffer (Li-Cor). Blots were probed with the appropriate primary antibodies overnight at 4°C in Odyssey blocking buffer supplemented with 0.1% Tween-20, followed by the secondary antibodies IRDye® 680RD donkey anti-mouse IgG or IRDye® 800CW donkey anti-rabbit IgG (Li-Cor) for 1 hour at room temperature. Fluorescent signals were imaged and quantified using Odyssey® CLx. Primary antibodies used were: HSP90 (BD Transduction Laboratories,

610419), phospho-p44/42 MAPK (ERK1/2) (Thr202/Tyr204) (Cell Signaling Technology, #4370), p44/42 MAPK (ERK1/2) (Cell Signaling Technology, #4695), phospho-MEK1/2 (Ser217/221) (Cell Signaling Technology, #9154), MEK1/2 (Cell Signaling Technology, #4694), phospho-AKT (Thr308) (Cell Signaling Technology, #13038), phospho-AKT (Ser473) (Cell Signaling Technology, #4060), AKT (Cell Signaling Technology, #4691 or #2920), phospho-EGF Receptor (Tyr992) (Cell Signaling Technology, #2235), phospho-EGF Receptor (Tyr1045) (Cell Signaling Technology, #2237), phospho-EGF Receptor (Tyr1068) (Cell Signaling Technology, #3777), EGF Receptor (Cell Signaling Technology, #4267), phospho-NF-kB p65 (Ser536) (Cell Signaling Technology, #3033), NF-κB p65 (D14E12) (Cell Signaling Technology, #8242), phospho-IκBα (Ser32) (Cell Signaling Technology, #2859), IκBα (L35A5) (Cell Signaling Technology, #4814), NanoLuc (Promega, N7000) and pan-RAS (Sigma, SAB1404011).

**Pull-downs:** Cells were lysed in ice-cold cell lysis buffer (Cell Signaling Technology) supplemented with 1 mM PMSF and Complete™ EDTA-free protease inhibitor cocktail (Roche) for 30 min with intermittent vortexing. Lysates were centrifuged at 18,000 g, 4°C for 15 min and supernatants were snap frozen in liquid nitrogen. Protein concentration was determined using the BCA protein assay kit (Pierce). 200 µg protein was mixed with 10 µl anti-HA magnetic beads (Cell Signaling Technology, #11846) in the presence/absence of peptides and incubated overnight at 4°C. The next day, beads were washed three times in ice-cold cell lysis buffer. After the final wash, bound proteins were eluted with 1X Bolt™ LDS sample buffer supplemented with NuPAGE® sample reducing agent and resolved using Western blot as described above. Primary antibodies used were: HA-tag (Cell Signaling Technology, #2367), b-RAF (Santa Cruz Biotechnology, sc-5284), c-RAF (Cell Signaling Technology, #12552).

**Traditional CETSA Assay:** Macrocyclic peptide cellular target engagement to $KRAS^{G12D}$ (GDP) were characterized using traditional western blot CETSA. AsPC-1 cells (Homozygous $KRAS^{G12D}$) were seeded in 6 well plates (0.25 million cells/2 ml in RPMI + 10% FBS). The following day, cells were treated with peptides dose dependently for 1hr and then trypsinized (0.25%) and pelleted after spinning at 300 rpm for 5 min. After a PBS wash, the cell pellets were resuspended with PBS (30uL/well/dose concentration) containing protease inhibitor (Nacalai Tesque) and transferred into 96 well PCR plate (Biorad) and then heated at 71°C for 3 min followed by cooling to 25°C for 3 min using eppendorf Mastercycler X50a PCR machine. Note that the melting temperature (ΔTm) for KRAS protein is previously determined to be 71°C [7]. After this step, the cells were frozen immediately in liquid nitrogen and stored at -80°C. The cells were then thawed and 30uL of 2X stock lysis buffer (100 mM HEPES (pH 7.5), 10mM beta-glycerophosphate, 0.2 mM Sodium orthovanadate, 20 mM MgCl2, 4mM TCEP, 0.05% of w/v Rapigest (Waters) and protease inhibitor (Nacalai Tesque)) were added to each PCR well. TCEP and Rapigest were prepared freshly. After the addition of lysis buffer, the cells were frozen and thawed three times in liquid nitrogen and were spun at 21000g for 20 min, 4°C. Supernatant were collected and transferred to new tubes. Protein concentration was determined using the BCA protein assay kit (Pierce). 20 to 50 µg of protein extract was separated on 4-12% Bis-Tris plus gels, transferred onto nitrocellulose membranes using the Trans-Blot® Turbo™ semi-dry system (Bio-rad), and blocked for 1 hour at room temperature with tris-buffered saline (TBS) Odyssey blocking buffer (Li-Cor). Blots were probed with KRAS specific antibody (Thermofisher, #703345) and SOD1 as a loading control (Cell signaling, #4266) overnight at 4°C in Odyssey blocking buffer supplemented with 0.1% Tween-20, followed by the secondary antibodies IRDye® 680RD donkey anti-mouse IgG or IRDye® 800CW donkey anti-rabbit IgG (Li-Cor) for 1 hour at room temperature. Fluorescent signals were imaged and quantified using Odyssey® CLx.

**Fluorescence imaging:** Cells were seeded in 96-well poly-D-lysine coated µCLEAR® plates (Greiner) and allowed to attach overnight. The next day, cells were treated with the indicated FAM-labelled peptides for 30 min. Nuclei were counterstained with Hoechst and dead cells were stained with TO-PRO™-3 Iodide. For the imaging of eGFP fusion proteins, AsPC-1 cells were transfected with the corresponding mRNAs for 6 hours before treatment with the indicated peptides for 1 hour. mRNA synthesis and transfection were performed as previously described [8]. Images were acquired using the Opera Phenix™ High Content Confocal Screening System under the 40X water immersion lenses and quantified using the integrated data analysis software.

**Cell Homogenate Stability and Metabolite Identification:** Stability of peptides towards intracellular proteases were evaluated using homogenized cells. Suspended THP1 cells at 1 million cells/mL were sonicated in bursts with a probe sonicator on ice until uniformly homogenized. The homogenate thus prepared was frozen and stored at -20°C until use. The peptides were incubated with the homogenate and loss of the peptide with increasing time was quantified using LC-MS/MS. Briefly, the test peptide was incubated with the homogenate at 1 µM in final volume of 50 µL at 37°C. At 0, 10, 30, 60 and 120 min, the reactions were quenched with 150 µL solution of 80% (v/v) Acetonitrile in Methanol containing a known internal standard peptide. The quenched samples were centrifuged at 4000 rpm for 5 min at 10°C and supernatants injected for bioanalysis by liquid chromatography combined with tandem-mass spectrometer. Pre-boiled homogenate (prepared by incubating homogenate for 30 min in boiling water) was also used as a negative control for each test peptide. The $k_e$ is the first-order rate constant describing the disappearance of parent drug in the incubation and obtained from regressing the initial slope (ke = -(slope)) of the natural log of the analyte area/ internal standard area (designated as C at an appointed time t) in C versus time (min) profile. The half-life ($t_{½}$) is calculated as:

$$t½ = \frac{0.693}{ke}$$

For metabolite identification, peptides were incubated in the homogenized cells similarly for 10 min and 30 min, and the high-resolution mass spectrometric analysis was performed, and metabolites assigned based on the mass to charge ratios.

**In Vitro Rat Mast Cell Degranulation and Histamine Release Assessment:** Peritoneal lavage was isolated from Wister-Han rats as described in literature. In brief, animals were anesthetized with xylene followed by an injection of about 10 mL of Tyrode buffer (12 mM $NaHCO_3$, 127 mM NaCl, 5 mM KCl, 0.5 mM $NaH_2PO_4$, 1 mM $MgCl_2$, 5 mM Glucose and 10 mM HEPES, pH 7.4) into the peritoneal cavity. After a gentle abdomen massage, peritoneal lavage was collected and centrifuged at 350 x g for 5 minutes. Cell pellets were resuspended in Tyrode buffer with 10% heat inactivated rat serum to reach a final concentration of 1 X $10^6$ cells/mL. Freshly isolated rat peritoneal cells were aliquoted to a 96 well tissue-culture plate (100 µL per well). Cells (1 X $10^5$ /mL) were incubated with titrating concentrations of various peptides for 30 minutes at 37°C. After incubation, the plate was centrifuged at 350 x g for 2 minutes at room temperature. Supernatant were collected and histamine concentration was measured by ELISA (Neogen, Lansing, MI) following vendor's instructions. Amount of histamine in the supernatant was presented as ng/mL. A detection of histamine release was defined as the histamine concentration in supernatant that was higher than the mean control values plus 3 standard deviations (typically derived from 10 vehicle-control treated samples from the same animal). The lowest concentration that a histamine release was observed for each peptide was summarized in table X (averaged from 2 rats).
